# STHD: probabilistic cell typing of single Spots in whole Transcriptome spatial data with High Definition

**DOI:** 10.1101/2024.06.20.599803

**Authors:** Chuhanwen Sun, Yi Zhang

## Abstract

Recent spatial transcriptomics (ST) technologies have enabled single- and sub-cellular resolution profiling of gene expression across the whole transcriptome. However, the transition to high-definition ST significantly increased data sparsity and dimensionality, posing computational challenges in identifying cell types, deciphering neighborhood structure, and detecting differential expression - all are crucial steps to study normal and disease ST samples. Here we present STHD, a novel machine learning method for probabilistic cell typing of single spots in whole-transcriptome, high-resolution ST data. Unlike the current binning-aggregation-deconvolution strategy, STHD directly models gene expression at single-spot level to infer cell type identities without cell segmentation or spot aggregation. STHD addresses sparsity by modeling count statistics, incorporating neighbor similarities, and leveraging reference single-cell RNA-seq data. We show in VisiumHD data that STHD accurately predicts cell type identities at single-spot level, which achieves precise segmentation of both global tissue architecture and local multicellular neighborhoods. The high-resolution labels facilitate various downstream analyses, including cell type-stratified bin aggregation, spatial compositional comparisons, and cell type-specific differential expression analyses. Moreover, STHD labels further reveal frontlines of inter-cell type interactions at immune hubs in cancer samples. STHD is scalable and generalizable across diverse samples, tissues, and diseases, facilitating genome-wide analyses in various spatial organization contexts. Overall, computational modeling of individual spots with STHD facilitates discoveries in cellular interactions and molecular mechanisms in whole-genome spatial technologies with high resolution. STHD is available at https://github.com/yi-zhang/STHD.

## Background

Spatial transcriptomics (ST) technologies have enabled gene expression profiling in the native spatial context of tissues. These technologies provide a systematic approach to profile cell type distribution, regional variations, and cellular neighborhoods across normal and disease samples. Current ST technologies are based on next-generation sequencing or in situ hybridization imaging, in short, seq-ST and image-ST(1). Often, seq-ST approaches cover towards whole-transcriptome with compromised resolution or depth (e.g. 10X Visium, Slide-seq(2), Stereo-seq(3)), whereas image-based approaches can provide sub-cellular resolution measurement of hundreds of genes in a designed panel (e.g. 10X Xenium, STARmap(4), MERFISH(5)). The technical tradeoff between resolution and throughput determined that seq-ST is generally suitable for hypothesis generation while image-ST for mechanism validation(6).

Emerging ST technologies have rapidly advanced with image-ST increasing gene coverage and seq-ST enhancing resolution to sub-cellular scale. Notable technologies that scale to whole-transcriptome and provide single- or sub-cellular resolution include HDST(7) (2um spot), Seq-Scope(8) (0.5-0.8um resolution), Stereo-seq(3) (spot size about 0.22um and center-to-center distance 0.5 or 0.715um), and Open-ST(3) (center-to-center distance 0.6um), *etc*. Recently, VisiumHD from 10X Genomics is further available for formalin-fixed paraffin-embedded (FFPE) samples to profile whole transcriptome in spots of size 2×2um, a sub-cellular size for most cell types and tissue contexts(9). Despite these technological advancements, the analysis of this data type presents significant computational challenges. The read coverage in each high-resolution spot is often highly sparse, and it is usually required to bin adjacent spots at a chosen size. For example, aggregation of 4×4 spots into 8×8um bins is usually recommended in VisiumHD sample processing. Similarly, typical Stereo-seq analyses aggregate data by binning 50 spots(3,10). The binning process computationally merges spots for sufficient read depth in downstream analytical tasks(11). However, this approach creates bins spanning multiple single cells, potentially mixing distinct cell types and thus compromising resolution. To address this, computational deconvolution of cell proportions is further needed, similar to the scenario in low-resolution seq-ST using tools like RCTD(12), CARD(13), Cell2Location(14), *etc*. Furthermore, the vast number of million spots or bins in high-resolution ST data poses additional challenges in computational efficiency.

We here developed STHD (probabilistic cell typing of single <u>S</u>pots in whole <u>T</u>ranscriptome spatial data with <u>H</u>igh <u>D</u>efinition), a machine learning method designed for cell typing of individual spots in high-resolution, whole-transcriptome spatial data. STHD reverses the traditional approach by first inferring cell type identities directly at the raw spot level, without requiring cell segmentation or bin aggregation. This is followed by cell type-specific bin aggregation as needed for downstream genome-wide analyses. To tackle high dimensionality and data sparsity, the STHD model employs a neighbor-augmented loss. It leverages cell type-specific gene expression from reference single-cell RNA-seq data, constructs a statistical model on spot gene counts, and employs regularization from neighbor similarity. The output includes cell type labels and probabilities for each individual spot. The STHD labels further facilitate multiple downstream analyses, including cell type-specific compositional comparison, cell type-specific differential expression, global tissue architecture segmentation, local cellular niche identification, cell type-augmented binning, and cell-cell interactions analyses. We demonstrated application of STHD to a colon cancer VisiumHD sample to uncover spatially heterogenous cancer cell expression programs and identifying immune infiltrate hubs with diverse immune functions and interactions. Moreover, we demonstrated that STHD is scalable and generalizable across different tissues, providing both local subcellular classification and global tissue segmentation.

## Results

### STHD efficiently models high-resolution spots using a machine learning model with neighbor regularization

An overview of STHD model is illustrated in **Fig.1a**. STHD infers latent cell type identities of each spot using a loss function that simultaneously optimizes two components: likelihood of spot gene counts and similarity with neighbor spots (**Fig.1a**, **Methods, Supplementary File 1**). The first part optimizes log likelihood of spot gene counts following Poisson distribution with the latent cell type variable, where parameters are based on normalized gene expression derived from a reference scRNA-seq dataset with cell type annotation from a matched disease or system. The second part denoises sparsity using cross entropy loss based on neighborhood similarity in cell type probabilities. The contribution from neighbors is controlled by the neighbor parameter β, which can be tuned with automatic stopping criterion to achieve balance between statistical likelihood and neighborhood similarity. For each high-resolution spot, STHD outputs cell type probabilities and labels based on Maximum a Posterior (MAP). The spot cell types enable multiple downstream analytical tasks, including cell type-stratified bin aggregation, cell type-specific compositional and differential gene expression analyses, and cell-cell interaction analyses. STHD implements fast optimization enabled by gradient derivation, numba(15), and Adam(16) optimizer (full derivation of loss gradient in **Supplementary File 1**). STHD also includes local low-count detection to filter cell and tissue gaps (**Methods, Supplementary Fig.1**). To be compatible with the millions of spots in one sample, we built the STHD pipeline that parallelizes patch-level inference followed by whole-sample integration (**Supplementary Fig.2**). The pipeline is thus adaptive to various computing resources without requiring GPU for optimization. Finally, to enable close examination of cellular neighborhoods, we developed STHDviewer for interactive and scalable visualization of spots in the entire sample (**Supplementary Fig.2**).

**Fig. 1.**
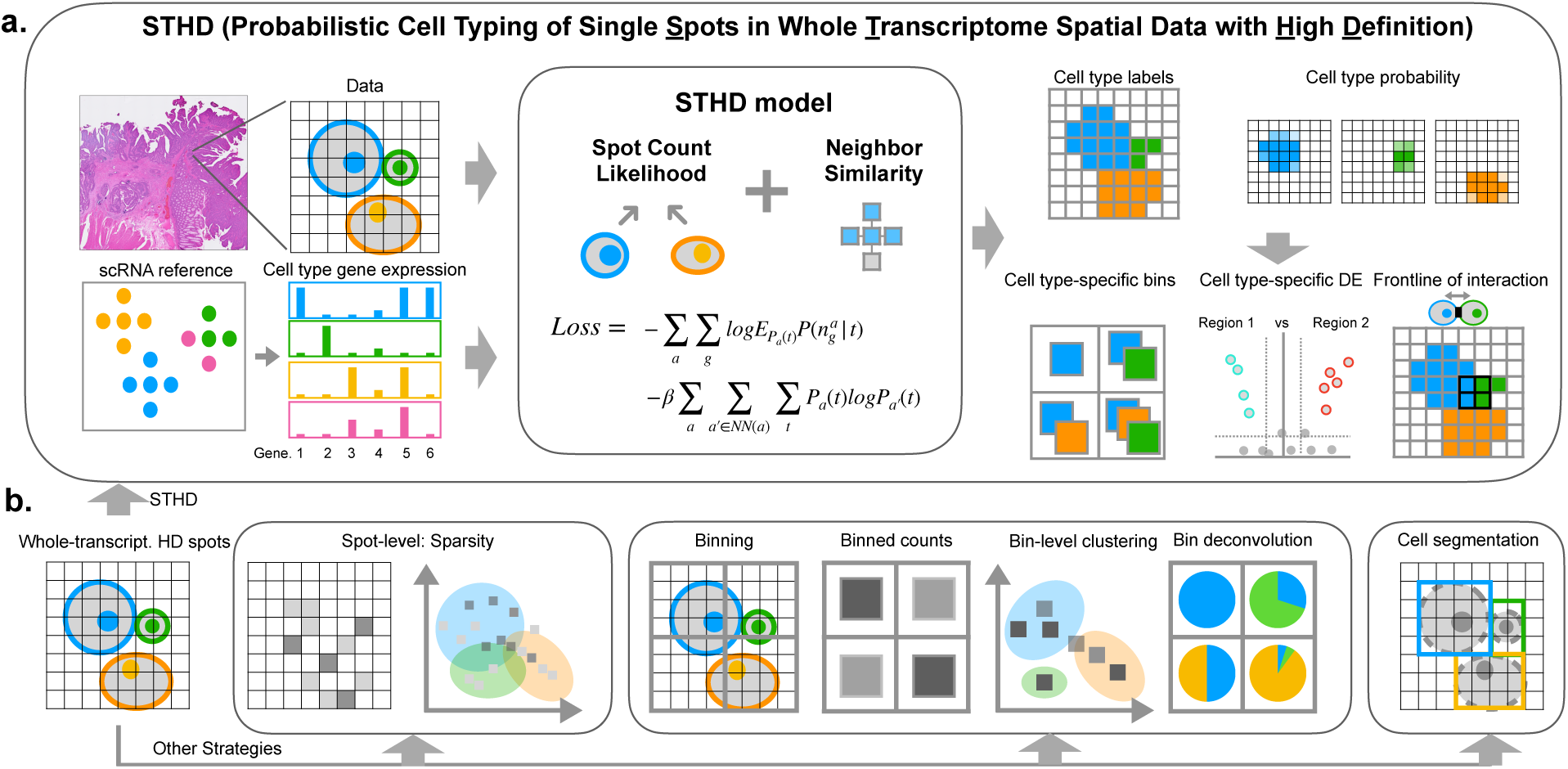
Overview of STHD model. **a.** STHD (probabilistic Cell Typing of Single <u>S</u>pots in Whole <u>T</u>ranscriptome Spatial Data with <u>H</u>igh <u>D</u>efinition) takes two inputs: whole-transcriptome spatial data of high resolution, and gene expression profiles by cell type from reference single-cell dataset. The STHD model is a machine learning method that infers latent cell type identity of individual spots by simultaneously modeling gene counts and neighbor similarity. STHD outputs cell type identities for each single spot in formats of cell type labels and posterior probabilities. The spot-level cell identities can then be used for multiple downstream analyses, including STHD-guided binning with cell type stratification, spatial comparison of cell type-specific compositional and differential expression analyses, and cell-cell interaction frontline analyses. **b.** Alternative approaches to analyze the HD datatype. Left, direct unsupervised clustering of spot-level gene expression is affected by sparsity. Middle, current binning-deconvolution approach increases coverage but compromise resolution. Right, ideal cell-level aggregation based on segmentation.

Different from current standards that start from binning, we propose in STHD that the analyses of whole-transcriptome, high-resolution ST data can begin with modeling the sub-cellular HD spots. This strategy offers advantages compared to alternative approaches. First, direct unsupervised clustering of spot-level gene expression results in false grouping of spots due to extreme sparsity (**Fig.1b**, left). Second, the current bin-aggregation approach can increase depth, but it mixes cell types and compromise resolution (**Fig.1b**, middle). Lastly, segmenting cell areas based on the hematoxylin and eosin (H&E) histological images may provide precise nucleus boundaries, but whole-cell segmentation has remained a challenging task(17) (**Fig.1b**, right).

### STHD infers cell type identities at the spot level, outperforms other inference tools, and guides cell type-specific binning

We first demonstrated STHD using the publicly available human VisiumHD sample from 10X Genomics, where a 6.5×6.5mm area of human colorectal tumor was profiled by 8,726,600 HD spots covering 18,085 genes. We constructed the normalized gene expression reference from an independent human colon cancer atlas study that profiled 43,113 genes from 370,115 single cells annotated into 98 fine-grained cell types from GSE178341(18) (**Supplementary Fig.3**). The reference gene profile construction step of STHD selected 4618 cell type-informative genes (**Methods, Supplementary Fig.3**). **Fig.2a** illustrated STHD label for a tumor-containing patch in the colorectal cancer sample containing 22,336 spots in a 300×300um area or 1100×1100 pixels in the full-resolution H&E image. First, we observed that STHD-predicted cell type labels collectively captured the compartmentalization of colon crypt-like structure with tumor-like epithelial cell type signatures (**Fig.2a**). In detail, stem-like proliferating cells were positioned inside, circled by the abundant enterocyte cells with scattered tumor stem and transit amplifying (TA)-like cells(18,19). The intercrypt regions were composed of abundant endothelial and fibroblast cells, with diverse immune cell populations such as plasma cells, monocytes, macrophages, dendritic cells, and T cells (**Fig.2a**). Interestingly, the cell type probabilities displayed high confidence for abundant epithelial spots as well as for rarer cells like macrophages and plasma IgG B cells (**Fig.2b**, **Supplementary Fig.4**). By default, spots with maximum posterior probability below 0.8 were assigned as ambiguous. The ambiguous spots mainly located at boundaries of different cell types and tissue structures and could represent cell type mixtures or cell gaps (**Fig.2a, b**). In comparison, firstly, clustering of spot-level gene expression vaguely reflected colon crypt structure with mingled cell identities (**Fig.2c, Supplementary Fig.5a**). This contradicts the expectation of consistent cell type labels within each cell, given that the raw spots (2×2um) are much smaller than a single cell (about 10um in diameter). Secondly, clustering of gene expression of bins aggregated from 4×4 spots (8×8um bins) within the patch resulted in less noise in crypt center and intercrypt regions (**Fig.2d** left**, Supplementary Fig.5b**). Moreover, similar bin grouping from the whole-sample clustering result could clearly distinguish the stemness center, enterocytes, and intercrypt regions, but the boundaries are assigned as one separated cluster that mixed in intercrypt immune cells (**Fig.2d** right). Further deconvolution on the aggregated bins using RCTD separated epithelial structure and immune cells but demonstrated a loss of spot-level resolution (**Fig.2e**).

**Fig. 2.**
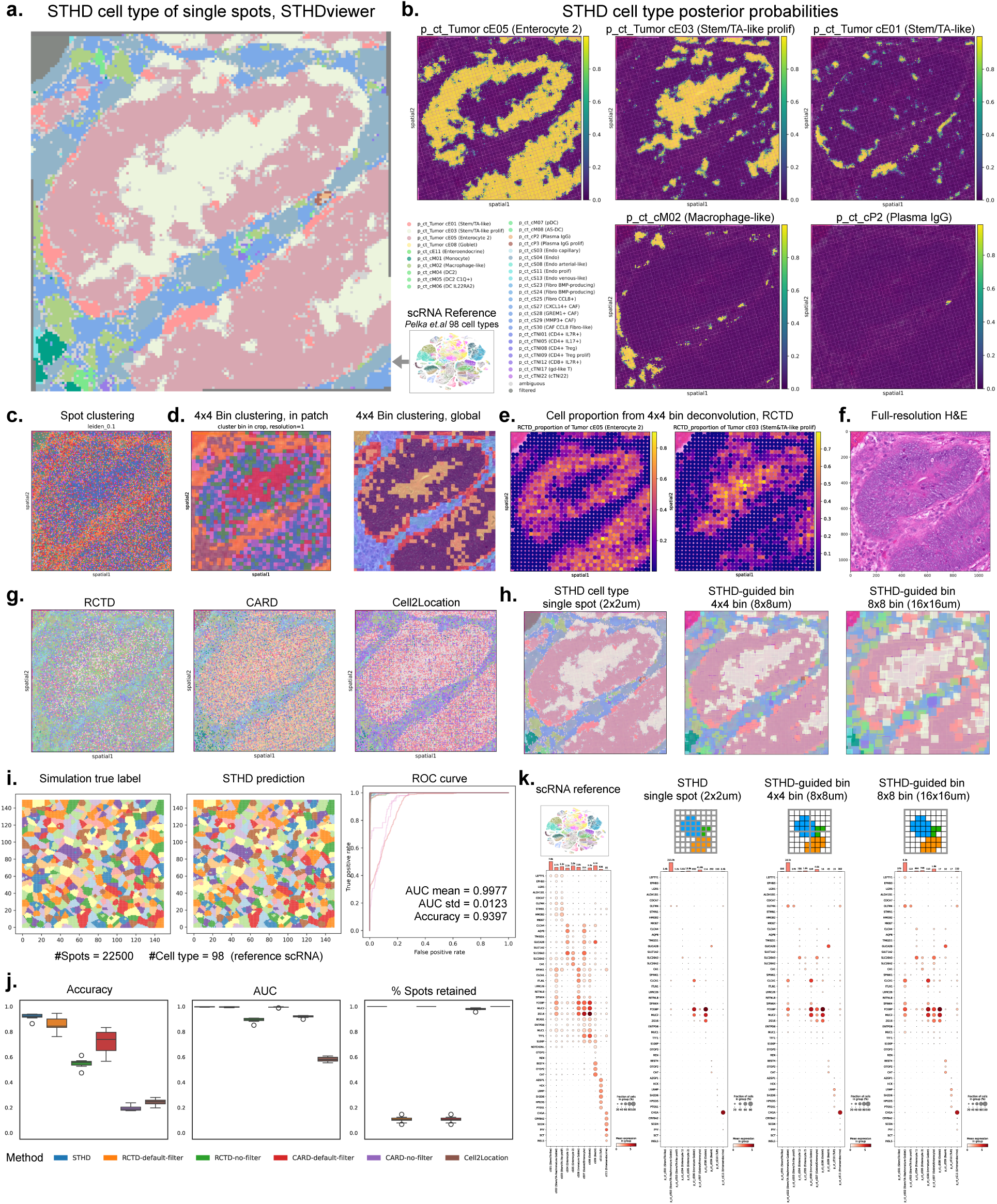
Spot-level predictions and benchmarking of a zoomed patch in a colon cancer VisiumHD sample. **a.** Example of STHD results of a patch from the human colon cancer VisiumHD sample, revealing tumor-like epithelial cells and intercryptic immune population, visualized using the interactive STHDviewer. **b.** Cell type posterior probabilities at spot-level for the same patch, visualized using Squidpy. Top row, three epithelial cell types; Bottom row, two immune cell types. **c.** A spatial plot of unsupervised leiden clustering on spot gene expression. **d.** Spatial plots of unsupervised leiden clustering of gene expression aggregated by bins in size of 4×4 spots. Left, clustering bins within patch, right, global cluster group of bins. **e.** Cell type proportion decomposed by RCTD on bins of size of 4×4 spots, demonstrating two tumor epithelial classes, enterocytes, and stem-like cells. **f.** The H&E histopathology image in full resolution of the same crop. **g.** Comparison to other cell typing and clustering methods adaptive to high-resolution spots. **h.** The STHD predicted labels at spot-level and STHD-guided binning in bins of 4×4 spots and 8×8 spots, visualized using Squidpy using the same colon cancer cell type colormap. **i.** Benchmarking with simulated high-resolution spatial data. Left, ground truth cell type labels; middle, STHD predicted cell type labels; right, receiver operating characteristic curve (ROC) curve for all cell types. **j.** Benchmarking against other methods for spot annotation in repeated spatial transcriptomics simulations. **k.** Expression dot plots for marker genes of normal epithelial cells, at spot level or STHD-guided bin level. Left to right: normal epithelial cell types in reference human colon cancer sample, normal epithelial cell types based on STHD predicted spots, normal epithelial cell types from STHD-guided bins of size 4×4 spots, normal epithelial cell types in STHD-guided bins of size 8×8 spots. The same cell type-specific gene markers from the original colon cancer study were used. ROC, Receiver operating characteristic. AUC, area under the curve.

Since high-resolution ST data lacks ground truth of spot labels, we evaluated the performance from aspects of histopathology, comparison to other methods, simulation, and marker gene expression. First, the spatial distribution of STHD cell type labels aligns with cell organization morphology based on H&E staining (**Fig.2f**). Second, we compared to cell type deconvolution on 8×8um bins by RCTD doublet mode or full mode using the same reference human colon cancer scRNA-seq reference (**Methods**). RCTD also resolved cell type proportions that match the colon crypt organization and the distribution of STHD labels for enterocytes and Stem/TA-like proliferating cells (**Fig.2e**, **Supplementary Fig.6a, b**). At the raw 2×2um spot level, we next tested computational methods that were able to label cell types by reference-based deconvolution like RCTD(12) and CARD(13), Cell2Location(14), Celloscope(20), Stereoscope(21), or by neighborhood-augmented clustering like Banksy(22). We noted that the common gene count filters in these tools need to be removed to avoid filtering of most spots; for example, 20436 of the 22336 spots are removed by RCTD default UMI cutoff 100 (**Supplementary Fig.6c**). At single-spot level, RCTD, CARD, Cell2Location, and Stereoscope can vaguely reveal the crypt structure while with highly inter-mingled cell types from different lineages (**Fig.2g**, **Supplementary Fig.7a-e**). The unsupervised Banksy efficiently separated epithelial crypt and intercrypt fibroblast/endothelial structures, but mixed the rarer immune cells inside the fibroblast/endothelial cluster and called the boundary as a separate cluster (**Supplementary Fig.7f**). Moreover, STHD has highest computational efficiency among all methods, which enables efficient inference on the entire sample (**Supplementary Fig.8**).

Since binned outputs are commonly used in VisiumHD, we noted that the STHD labels at the subcellular 2×2um resolution can further guide cell type-specific binning in cell-size resolution, by aggregating adjacent spots of the same cell type, thereby improving gene coverage while preventing cell type mixing. With spot-level cell types, an appropriate bin size can be chosen, allowing separation of distinct cell types within the same bin areas for cell type-specific gene expression profiles. **Fig.2h** showcased STHD-guided binning with two commonly used bin sizes, 8×8um (single-cell size) and 16×16um; each cell type-specific bin was positioned at centroid of the covered spots of corresponding cell type. Thus, tumor epithelial cells and intercrypt endothelial, fibroblasts and immune cells were clearly separated on the STHD-guided binning spatial maps (**Fig.2h**, **Supplementary Fig.5c, d**).

### STHD achieves high performance in benchmark simulations and preserves cell type-specific expression profiles

To quantitatively assess performance of STHD, we simulated comprehensive whole-transcriptome, high-resolution ST data *in silico* (**Methods, Supplementary Fig.9**). As shown in **Fig.2i**, we simulated 500 cells with heterogeneous cell sizes and cell types on 22,500 spots, each with a true cell type label and gene counts following expression levels from the same human colon cancer reference. STHD achieved average area under the curve (AUC) 0.9977 with overall accuracy 93.97% for this multi-class classification task with 98 categories (**Fig.2i, Supplementary Fig.9**). STHD outperformed other popular spot-level cell type annotation approaches when tested in repeated simulations, achieving highest accuracy, AUC, and percentage of retained spots (**Methods, Fig.2j**). Next, we systematically investigated cell type-specific expression of marker genes based on STHD prediction across the entire colon cancer sample. Despite spot sparsity (**Supplementary Fig.5e, Supplementary Fig.3d**), the STHD labels displayed similar cell type-specific expression patterns of marker genes as in scRNA-seq reference data in normal epithelial cells (**Fig.2k**, left two panels, **Supplementary Fig.10a**) and all other cell lineages: tumor epithelial cells (**Supplementary Fig.10b**), myeloid cells (**Supplementary Fig.11a**), B and plasma cells (**Supplementary Fig.11b**), CD8T, CD4T, and other T cells (**Supplementary Fig.12, Supplementary Fig.13a**), fibroblasts (**Supplementary Fig.13b**), and endothelial cells and pericytes (**Supplementary Fig.14**). STHD-guided binning effectively increased marker gene coverage and meanwhile retained cell-type specificity, shown in two example bin sizes: 8×8um bins (**Fig.2k**, third panel, **Supplementary Figs.10-14**) and 16×16um bins (**Fig.2k**, fourth panel, **Supplementary Figs.10-14**). We also observed that the traditional binning approach that aggregated 8×8um bins yielded 311,396 bins with single cell type and 169,188 bins with more than one cell type, creating one third of 8×8um grid bins with mixed cell type signals.

### STHD enables scalable spatial inference and interactive exploration of cellular niches and immune hubs, facilitating genome-wide and cell type-specific spatial comparisons

The STHD pipeline enabled scalable inference across the entire ST sample through parallelized inference of spatial patches followed by whole-sample integration (**Supplementary Fig.2c**). The parameter tuning was thus performed for the colon cancer VisiumHD sample to balance between statistical likelihood and neighborhood similarity (**Supplementary Fig.15**). To facilitate close examination of spot identities and scalable investigation of cellular neighborhoods, we developed STHDviewer, an interactive and scalable visualization functionality capable of rendering millions of spots in a webpage (**Fig.3a, Supplementary Fig.16**). As demonstrated in **Fig.3a**, STHD prediction automatically achieved precise segmentation of global tissue architecture and local multicellular neighborhoods. Specifically, fibroblasts separated regions of normal-like area with colon gland morphology (named Epi-Normal-like, **Supplementary Fig.16-17**) and tumor epithelial areas at the margins (Epi-Tumor-left, Epi-Tumor-top, Epi-Tumor-right) compared to tumor in deposit (Epi-Tumor-inside)(23,24) (**Fig.3b**). Across tumor regions of interest (ROI), STHD labels delineated compositional heterogeneity of cell lineages and types (**Fig.3c**). Notably, spatial heterogeneity of immune cell populations was also observed; for instance, the Epi-Tumor-inside region contained macrophages(25,26) but few T cells compared to tumor regions at the margin, indicating distinct “hot” verses “cold” T cell infiltration spatial patterns within the same sample (**Fig.3c** top, **Supplementary Fig.18**).

**Fig. 3.**
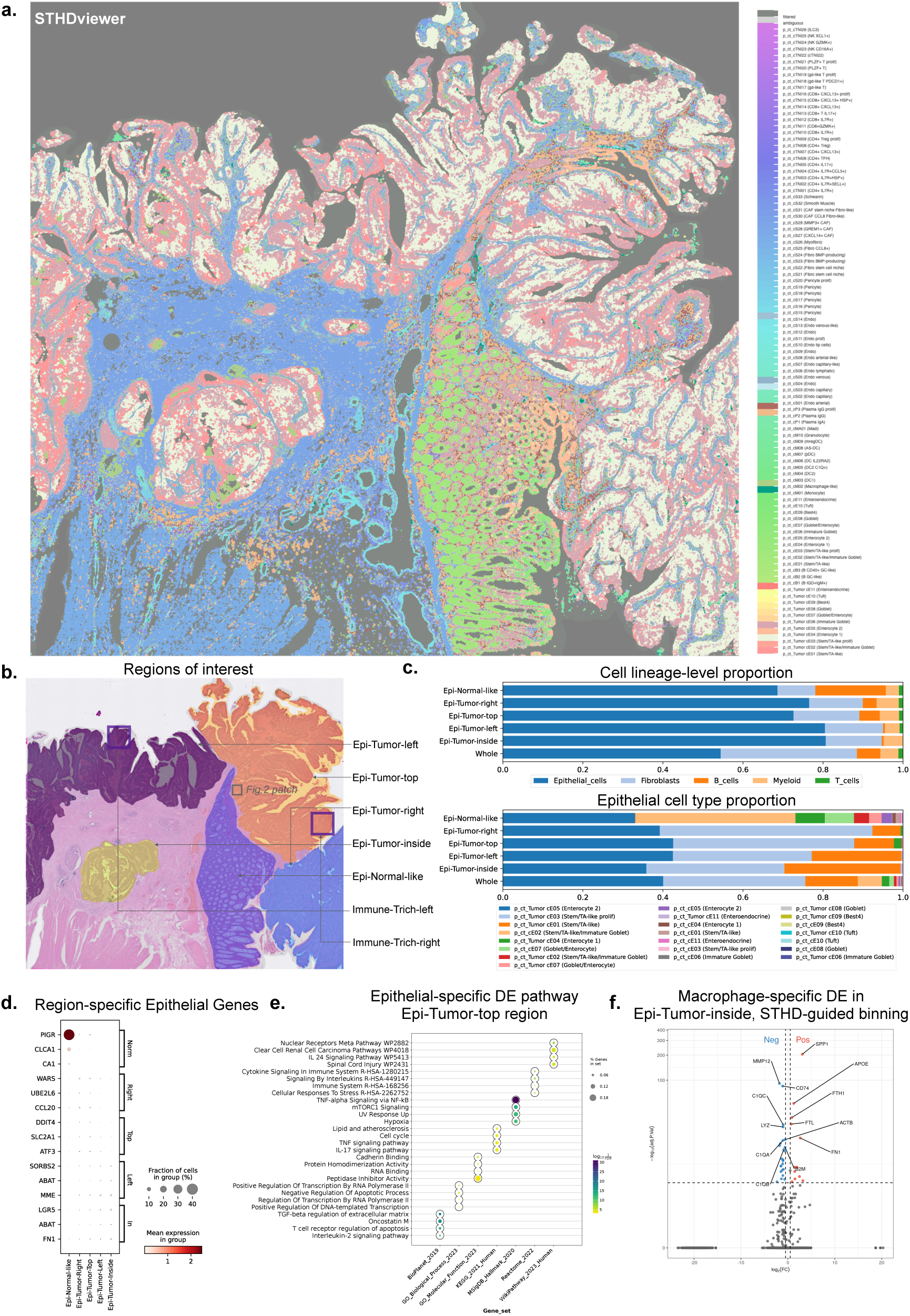
STHD prediction facilitates segmentation of global structures and cell type-specific differential analyses across spatial regions. **a.** Interactive spatial visualization by STHDviewer of predicted cell type labels for spots in the entire human colon cancer VisiumHD sample. **b.** Annotation of regions of interest, separating epithelial regions and two T cell-infiltrated regions. **c.** Cell type proportion bar plots at cell lineage and cell type level for epithelial regions and whole map. Upper: Proportions of major cell lineages including epithelial cells, fibroblasts, B cells, myeloid cells, and T cells. Bottom: Proportions of finer cell types of epithelial lineage. **d.** Spot-level gene expression for epithelial-specific differential genes across regions. Top three genes by log fold change were shown. **e.** Pathway enrichment of epithelial cell-specific genes differentially expressed in the Epi-Tumor-top region. **f.** Macrophage-specific differential genes for the Epi-Tumor-inside region, where differential analyses were performed at bin level of size 4×4 spots with cell type stratification from STHD. DE, differential expression.

Besides cell type compositional comparisons, STHD facilitates genome-wide and cell type-specific differential expression (DE) analyses across spatial regions. Comparing different tumor ROI, epithelial cells in Epi-Tumor-inside expressed highest stemness gene *LGR5*(27), whereas epithelial cells in Epi-Tumor-top highly express *CCL20*(28) and enrich pathways like TNF-alpha signaling (**Fig.3d,e, Supplementary Fig.19-20**). Since the Epi-Tumor-inside region enrich macrophages, we compared differentially expressed genes specific to macrophage spots and observed that macrophage infiltrating inside tumor expressed higher extracellular matrix gene *SPP1* and lower metalloelastase *MMP12*, cytokine *CXCL8*, and antigen presentation gene *B2M*, indicating that macrophage states segregated spatially in the same tumor(26,29,30)(**Fig.3f**, **Supplementary Fig.21**). The cell type-specific DE analyses can also be performed on the level of STHD-guided bins with higher depth, which resulted in a higher number of significant DE genes with well correlated effects (**Fig.3f**, **Supplementary Fig.20-21**).

We next demonstrated STHD in analyzing the less abundant immune cells after using STHDviewer to identify immune cell hubs at different locations in the colon cancer VisiumHD sample. An advantage of STHD is that rarer cell types and fine-grained cellular neighborhoods are maintained on top of the denoising effect for those abundant cell types forming continuous tissue architecture; an example of immune-endothelial cell interactions was demonstrated in **Supplementary Fig.22**. **Fig.4a** highlighted two T cell-rich areas marked in **Fig.3b**. The Immune-Trich-left region contained CD4T and CD8T cells with abundant macrophages and C1Q-high dendritic cells (DC)(31), while the Immune-Trich-right area contained CD4T, CD8T cells, mature DCs enriched in immunoregulatory molecules (mregDC)(32), and Plasma IgG B cells (**Fig.4b**). In neighborhood enrichment analyses, each area displayed unique co-occurrence patterns among immune cell types; for example, T cells in Immune-Trich-left area frequently interacted with macrophage and DCs, while T cells in Immune-Trich-right area co-occurred with DCs (**Fig.4a,c**). Utilizing cell type-specific DE analysis, we observed that T cells in the Immune-Trich-left region expresses higher cytotoxicity genes like GZMB comparing to T cells in the Immune-Trich-right region, which can be associated with the immunosuppressive function of mregDCs in tumor(33–35) (**Supplementary Fig.23**).

**Fig. 4.**
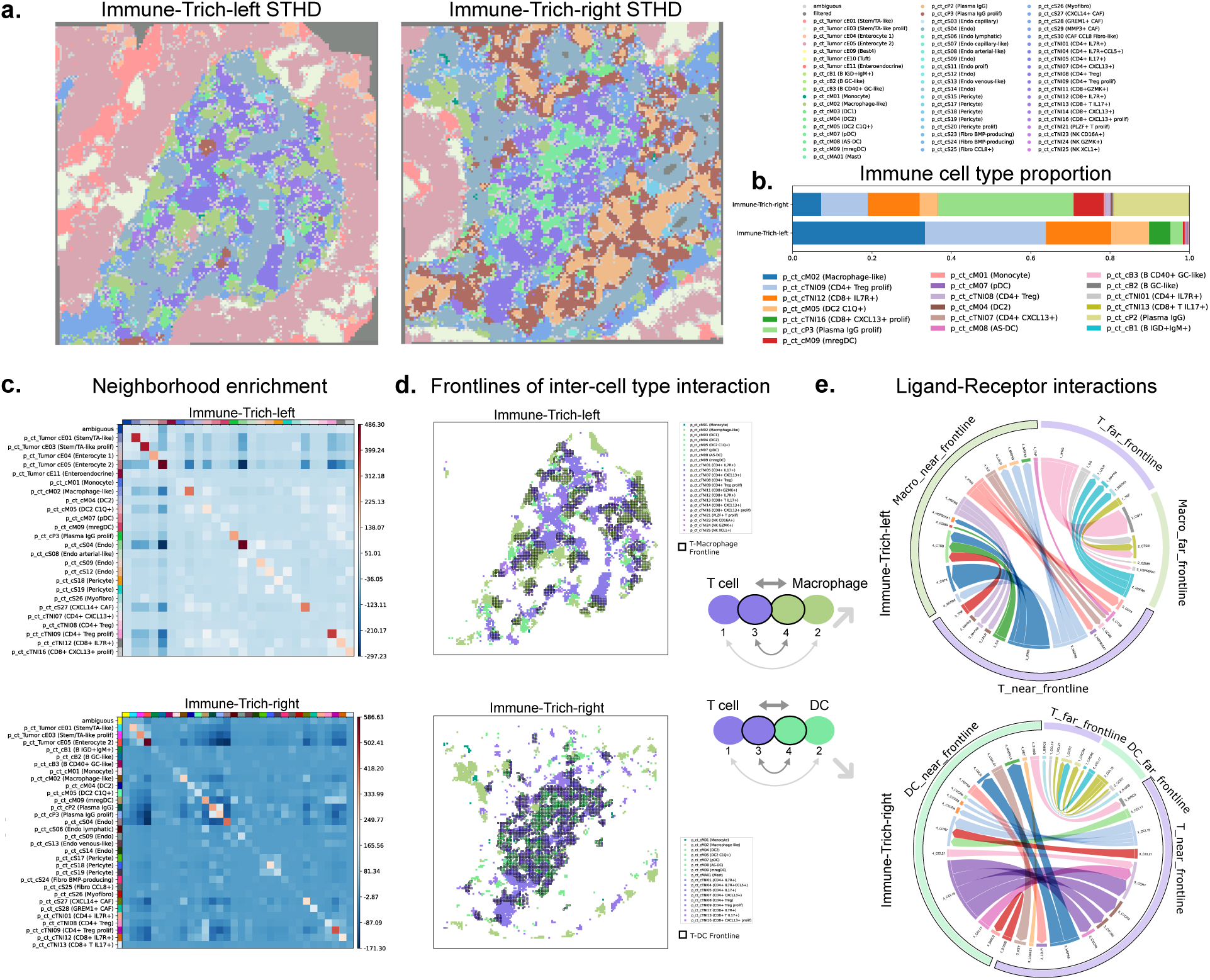
STHD enabled analyses of local neighborhoods for cellular interactions in tumor-infiltrating immune hubs. **a.** STHD cell type labels for two T cell-rich regions in the colon cancer sample. **b.** Proportions of immune cell types in the two immune regions. **c.** Cell neighborhood interaction patterns of the two T cell-rich regions showing neighborhood enrichment z-scores where spots within 6 spots are considered neighbors. **d.** T cell and myeloid cells in the two immune-rich regions, with frontlines of inter-cell type interactions. Top, T cell-macrophage frontline in Immune-Trich-left region; Bottom, T cell-dendritic cell frontline in Immune-Trich-right region. Considering a pair of cell types, spots within 4 spots (8um) from the inter-cell type interaction location are highlighted as near-frontline spots, with the rest as far-frontline spots. **e.** The ligand-receptor communications at near-frontline spots compared to far-frontline communication scores of the same cell type pair as background, where the ligand-receptor interaction scores were calculated by squidpy.ligrec. Top 10 ligand-receptor pairs different between near-frontline and far-frontline communications were visualized where width represents average expression level of ligand receptor pairs. Spots are classified into four groups 1,2,3,4 based on their cell type and near- or far-frontline categories.

### STHD identifies precise cell-cell interactions frontlines in immune hubs and reveals direct ligand-receptor communication activities

Cell-cell communications between cell types in disease microenvironment could drive disease progression like tumorigenesis(36,37). Molecular interactions between cells types in single-cell transcriptomics analyses are traditionally inferred by evaluating averaged expression of communicating gene pairs, which lacks support on direct and physical interactions(38). With the sub-cellular resolution spots labeled, STHD can pinpoint the precise frontlines of inter-cell type interactions. As shown in **Fig.4d**, we highlighted the frontlines of T cell-macrophage interaction and T cell-DC cell interaction in each of the T cell-rich regions. We hypothesized that cell-cell communications mediated by ligand-receptor signaling near-frontline could be stronger than that in far-frontline spots. For a given cell type pair, we identified spots at boundary of two cell types based on posterior probabilities and considered STHD-labeled spots within 4 spots (comparable to single cell size) in distance as near-frontline spots (**Methods, Fig.4d**). We calculated ligand-receptor interaction scores (LRIS)^21,22^ between the near-frontline spots and compared with background interaction scores from far-frontline spots, conditioning on the same cell type pair. As shown in **Fig.4e**, near-frontline spots (grouped as 3,4) harbored higher LRIS activities compared to the far-frontline spots (grouped as 1,2) in both cell type pairs of T-Macrophage (**Fig.4d,e** top) and T-DC interactions (**Fig.4d,e** bottom). Specifically, the T-Macrophage communications in Immune-Trich-left were largely driven by *CD74*-*IFNG* interaction, while the T-DC communications in Immune-Trich-right were driven by *CCL17*/*CCL19*-*CCR7* interactions (**Fig.3k**), which indicated the immunoregulatory effect of mature DCs on T cells(39).

### Multi-sample analyses of human colon cancer identified sample-specific immune enrichment patterns

As data accumulate for whole-genome spatial transcriptomics with high-definition, we next extended STHD to additional VisiumHD samples. Oliveira et al. recently profiled multiple colorectal cancer samples and adjacent normal colon tissues using VisiumHD(9). We applied STHD to analyze two cancer samples (P1, P5) and one adjacent normal colon sample (P3), in addition to the previously released sample (P2) included in this study. **Fig.5a** illustrated STHD-inferred cell type identities for colon cancer sample P1 containing 8,115,201 spots at 2×2um spot resolution. The colon cancer sample P1 exhibits a similar tumor organization to the colon cancer sample P2, with segmented tumor regions containing colon cancer epithelial cells interspersed with fibroblasts, and adjacent colon tissue enriched with goblet cells. Notably, the colon cancer sample P1 features a prominent tertiary lymphoid structure (TLS), characterized by abundant germinal center B cells and surrounding T cells (**Fig.5b** top). The TLS-like regions was also validated by B cell markers (*CD19*, *MS4A1*) and the TLS-associated T cell cytokine *CXCL13* (**Fig.5b** bottom)(40). In contrast, other colon cancer samples P2 and P5 lack TLS-like structures (**Fig.2, Fig.5c** left). STHD also provided detailed insights into the normal colon tissue from sample P3, revealing goblet cells, an adjacent lymphoid structure, and a myofibroblast region (**Fig.5c** right). Overall, STHD captures heterogenous cellular compositions across multiple samples highlighting distinct immune cell profiles (**Fig.5d**).

**Fig. 5.**
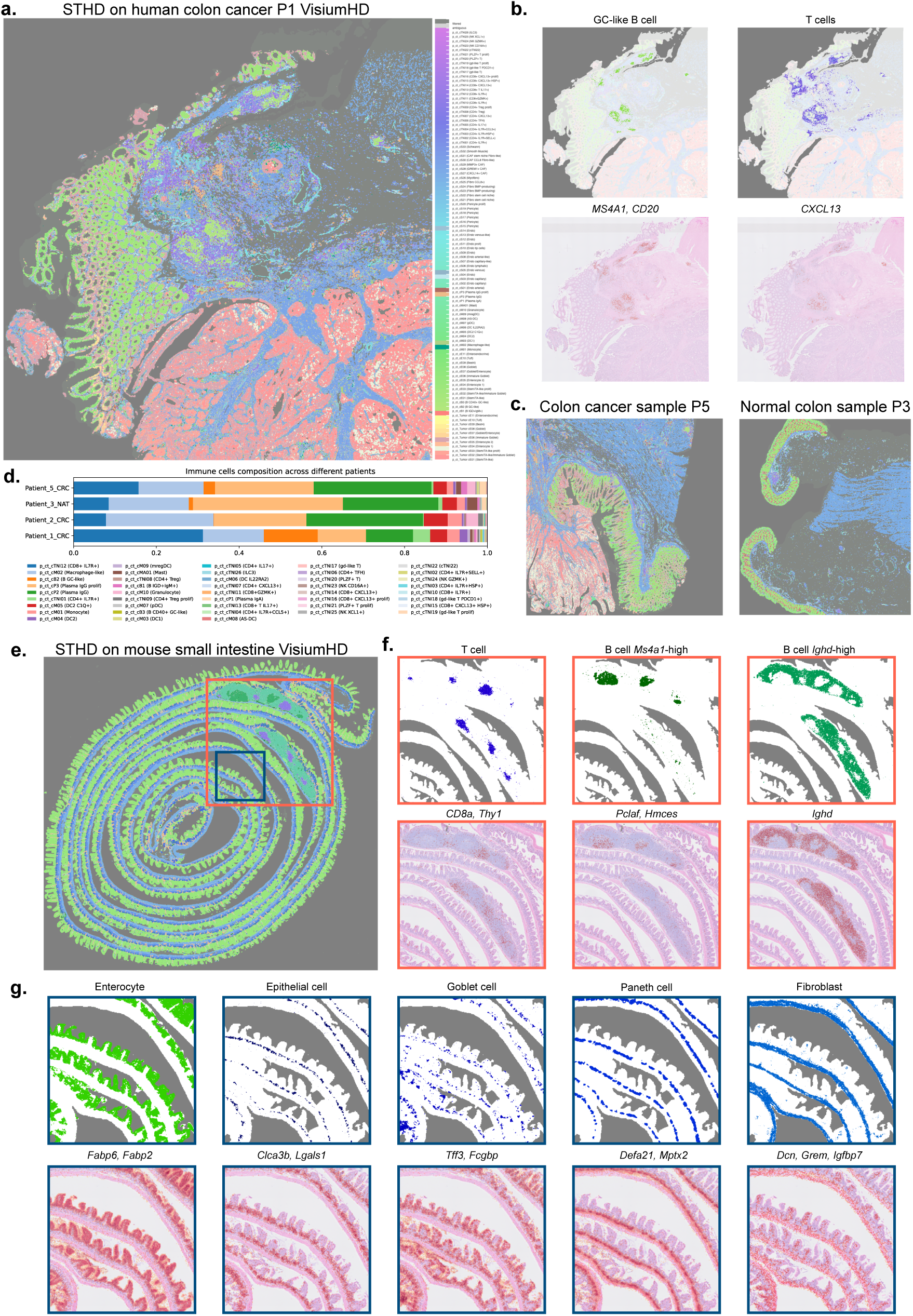
STHD is scalable to other samples, revealing subcellular cell type identities and tissue structures. **a.** STHD spot-level inference of a human colon cancer Visium HD sample showing tumor, colon tissue, and enriched lymphoid structure. **b.** Left, spots (2×2um) labeled as germinal center B cells, with B cell marker expression aggregated per 8×8um bins. Right, spots (2×2um) labeled as T cells, and expression of TLS-specific T cell cytokine *CXCL13* by 8×8um bins. Marker expression plots are illustrated by loupe browser. **c.** Spot-level inference of human colon VisiumHD sample P5 (cancer) and P3 (normal adjacent tissue). **d.** Immune cell-specific composition across all four human colon Visium HD samples. **e.** STHD results for the mouse small intestine VisiumHD sample, with two zoomed-in regions highlighted. **f.** Spots with immune cell identities in the lymph node-like region, showing T cell, *Ms4a1*+ B cell and *Ighd*+ B cells. Top, STHD labels in 2×2um spots, Bottom, marker gene expression in 8×8um bins. **g.** Spots of intestine cells forming layered structure, including intestine enterocyte cell, epithelial cell, goblet cell, Paneth cell, and fibroblasts. Top, STHD labels in 2×2um spots, Bottom, marker gene expression in 8×8um bins. GC, germinal center. CRC, colorectal cancer. NAT, normal adjacent tissue.

### STHD is generalizable to diverse samples across tissues, diseases, and organisms

We then extended the application of STHD to normal tissues from both human and mouse model systems. Using the STHD pipeline with default parameters, we analyzed three VisiumHD samples from 10X Genomics including human pancreas, mouse small intestine, and mouse brain (**Methods**). For the adult human pancreas sample, we constructed cell type-specific gene expression estimation using reference single-cell RNA-seq data from Baron et al.(41), which included 12,000 pancreatic cells from 4 adult donors profiled by inDrop and annotated 14 cell types. STHD inference, based on this reference, demonstrated the pancreas’s abundant acinar cells and embedded pancreatic islets - organized cell structure harboring endocrine cells (**Supplementary Fig.24a**, left). We observed that marker gene expression levels matched expected patterns for various cell types, such as beta cells (*IAPP*, *INS*), alpha cells (*GCG*, *TTR*), epsilon cells (*GHRL*), and pancreatic stellate cells (*RGS5*) (**Supplementary Fig.24a**, right)(41). For the mouse small intestine, we utilized a reference scRNA-seq profile from the Mouse Cell Atlas 2.0(42) (**Methods**). STHD inference of spot identities collectively reveals the tissue structure of the small intestine (**Fig.5e**). Specifically, we highlighted the mesenteric lymph node with immune cells and the layered intestine-epithelium organizing enterocytes, epithelial cells, goblet cells, Paneth cells, and fibroblasts (**Fig.5f,g** top row), with matched expression of cell type-specific gene markers(42) observed at 8×8um bin-aggregated level (**Fig.5f,g** bottom row). Finally, we mapped the mouse brain VisiumHD data to scRNA-seq data based on Allen Brain Atlas(43) (**Methods**). STHD revealed brain layer-specific cell types for each high-resolution spot and differentiated known cortical layers and hippocampus regions (**Supplementary Fig.24b**). Overall, we observe a high degree of spatial correspondence to tissue structures collectively by STHD spot-level labels, demonstrating STHD’s generalizability across diverse tissues and disease samples.

## Discussion

As spatial transcriptomics technologies advance towards whole-genome coverage at single- and sub-cellular resolution, the development of computational methods that offer both local precision and global scalability is crucial for enabling comprehensive genome-wide analyses across spatial regions, samples, and conditions. However, the enhanced dimension and resolution introduces challenges such as data sparsity. To address this issue, existing analytical approaches often rely on hard aggregation of adjacent spots, which compromises resolution and leads to the mixing of distinct cell types. We presented STHD, a machine learning method designed to model cell types at the single-spot level for high-definition, genome-wide spatial transcriptomics data. Unlike traditional methods, STHD achieves soft aggregation of cell type-specific gene expression information contributed by neighboring spots, effectively addressing the computational difficulties posed by sparse data through a neighbor-augmented Expectation Maximization (EM) model. Our benchmarking simulations and analyses of recently available public VisiumHD samples demonstrate STHD’s accuracy, precision, and efficiency. With genome-wide gene expression profiles, STHD’s spot-level cell type labels enabled a range of downstream analyses, including the study of global tissue architecture, local cellular neighborhoods, cell type-specific differential expression, cell type-augmented binning, and cell-cell interactions. The STHD pipeline and viewer are implemented to provid efficient modeling and interactive examination, scalable to the entire sample containing millions of spots. Moreover, we have shown STHD’s generalizability across different samples, tissues, and disease contexts.

A tradeoff has existed in ST technologies between resolution and throughput. For seq-ST technologies, which typically cover a high number of genes, the limitation in spot resolution often results in the mixing of gene expression signals from different cell types within a single spot (e.g. Visium’s 55um spot diameter). Previous computational efforts have aimed to estimate cell type proportions using spot-level gene expression decomposition, such as RCTD(12), Cell2Location(14), CARD(13), *etc*. As seq-ST technology progresses towards subcellular spot size with genome-wide coverage, we adopted a different assumption: most spots could be attributed to a single cell type. This assumption reduces model complexity and allows for estimation based on sparse readouts. Notably, STHD’s efficient optimization enabled analyses of an entire VisiumHD sample with millions of spots using around ten CPU hours without requirement GPU resource. In both normal and tumor tissue samples, we observed that this assumption held for most spots, except for a subset classified as ambiguous. STHD detected these ambiguous spots based on the multinomial cell type probability vector, where lack of a dominant cell type (default posterior cutoff 0.8) indicates ambiguity. Comparison to histology reveal that these ambiguous spots often correspond to regions like cell gaps, certain non-nucleus cytoplasm areas, and mostly inter-cell type boundaries. Consequently, these cell type boundary spots can be utilized in identifying inter-cell type transitions at the highest possible resolution; also, STHD’s estimated cell type probabilities could implicate spot-level compositions.

The data sparsity in gene counts presents a significant challenge in subcellular spot modeling. For instance, in the colon cancer sample P2, the average count depth of a 10X VisiumHD spot was 1/262 of that of a single cell from 10X scRNA-seq. Consequently, this sparsity often results in the exclusion of most spots in existing spot-level methods or generation of noisy cell type labels when each HD spot was treated independently. To tackle sparsity, STHD incorporates a soft regularization of the loss function based on cross-entropy among neighboring spots’ cell type probability vectors. This approach leverages the neighbor similarities in spatial data, as seen in previous methods like Banksy, which utilized Gabor filter to incorporate structured neighborhood contribution in unsupervised tasks like tissue segmentation and spatial clustering(22). Our results show that STHD labels not only captures the continuous zonation of those abundant cell types but also identifies rarer cell types, such as immune cells in tissue and tumor samples. The recovered continuity in the spot identities further enabled cell type-guided binning, which maintains cell type-specific marker expression and enhances gene coverage for downstream differential expression analyses. In a recent study, Bin2Cell also avoids traditional binning methods but operates on the raw VisiumHD spots by pooling spots overlapping the same nucleus identified from H&E image(44). In another recent study, TopACT also models neighbor aggregation, where the range of multiscale aggregation was optimized for cell type classification(45). Similar to the neighbor parameters in Banksy and multiscale design in TopACT, STHD allows for tunning of neighborhood aggregation effect: the β parameter and the iteration step parameter control the loss contribution from neighbors, and the neighbor size parameter can control the range of spatial similarity aggregation, making it adaptable to various cell sizes and tissue types.

Despite the advancements, several challenges remain in analyzing high-dimensional data with subcellular resolution. First, the lack of group truth for spot-level cell identities in tissue ST samples complicates the evaluation of method performance. Our spatial simulations, which created spots with true cell type labels matching scRNA atlases, provide a starting point for such evaluations, but future integration with other spatial technologies (e.g., image-ST or proteomics) could offer additional ground truth data. The STHD model is also generalizable to other high-resolution, high-dimensional spatial data platforms like Stereo-seq and Xenium 5K, *etc*. Second, the availability and quality of the reference scRNA-seq data can limit the applicability of STHD, as it relies on these references for cell type identification but does not discover new cell types, a common feature also existed for spot deconvolution methods. In cases where detailed and cell type annotation are lacking or cell type profiles are highly correlated, STHD label can provide an initial labeling, STHD can still provide initial labeling, allowing for further discovery of transcriptional variations within each cell type through clustering of STHD-guided bins. We anticipate expanding of STHD’s capabilities for semi-supervised cell type identification. Third, the issue of molecular diffusion has been found to exist across ST methods(46), which may interfere with STHD prediction particularly in spots with weak marker gene signals. Computational tools to correct spot diffusion, such as SpotClean(47) for low-resolution data, have been developed, but there remains a need for methods compatible with millions of sparse spots. Additionally, molecular diffusion could also blur the tissue boundaries, as observed in some regions outside H&E images containing gene counts. This also imply that integrating histopathology properties can be informative in modeling the high-resolution spatial transcriptomics sample, as done in recent studies like iStar, which infers spatial gene expression from H&E images(48).

Looking ahead, we believe that the combination of individual subcellular spot modeling, neighborhood information sharing, and multi-modality neighbor information incorporation, including H&E images, will be key to overcoming the resolution-throughput tradeoff in emerging spatial transcriptomics technologies. This rationale fully harnesses the potential of recent advancements in spatial technology and enables the discovery of spatially organized cellular heterogeneity in samples. The algorithm design of STHD is thus generalizable in other platforms of high-dimensional spatial transcriptomics profiling with subcellular resolution. In summary, STHD offers probabilistic subcellular spot-level predictions that facilitate precise and scalable analyses, with a software toolkit that is generalizable across various tissues and diseases, providing new insights into cellular and molecular landscapes.

## Conclusions

We have presented STHD, a novel machine learning method for probabilistic cell typing of single spots in whole-transcriptome, high-resolution ST data. Unlike the current binning-aggregation-deconvolution strategy, STHD directly models gene expression at single-spot level to infer cell type identities without cell segmentation or spot aggregation. STHD addresses sparsity by modeling count statistics, incorporating neighbor similarities, and leveraging reference single-cell RNA-seq data. We show that STHD accurately predicts cell type identities at single-spot level in simulated benchmarking and biological spatial samples, where STHD achieves precise segmentation of both global tissue architecture and local multicellular neighborhoods. The high-resolution STHD labels facilitate various downstream analyses, including cell type-stratified bin aggregation, spatial compositional comparisons, and cell type-specific differential expression analyses. Moreover, STHD labels further reveal frontlines of inter-cell type interactions at immune hubs in cancer samples. Finally, STHD is efficient, scalable, and generalizable across diverse samples, tissues, and diseases. Overall, STHD will facilitate biomarker and mechanistic discoveries in spatial heterogeneity, cellular interactions, and molecular mechanisms in various tissue and disease contexts.

## Methods

### STHD algorithm

The STHD model is a machine learning method that combine modeling of per-spot gene counts based on latent cell type identities with regularizing similarity among neighboring spots in terms of cell type probability distribution. The overall loss function ℒ composes of negative log likelihood −*LL* that models gene count of all spots and the cross entropy loss *CE* measuring similarity of neighboring spots. The amount of contribution from neighbors is controlled by the parameter β. The cell type identities at every single spot are thus estimated by Expectation Maximization.

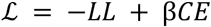

For the negative log likelihood part, we model counts of each gene at each spot to follow Poisson distribution. Let the normalized average gene expression level for gene *g* within the latent cell type *t* is 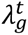, read depth of spot *a* is *d*, and spot *a*’s probability of belonging to cell type *t* is *P_a_*(*t*). Thus, the count of gene *g* at spot 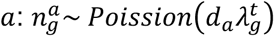.

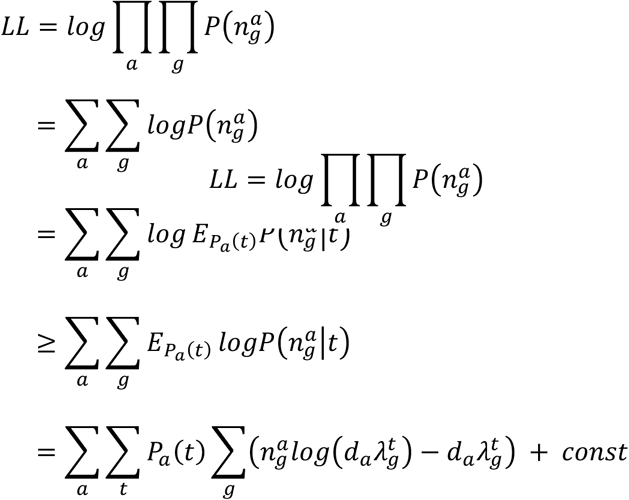

Considering high sparsity of single spots, the second part of the loss function is related to cross entropy loss between spot *a* and nearest neighboring spots *a′* ∈ *NN*(*a*) in terms of distribution of cell type probability.

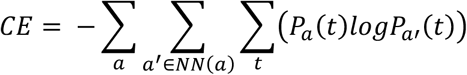

Note that one constraint is ∑*_t_ P_a_*(*t*) = 1. In calculation, we use the following for inherent incorporation of constraint: 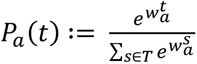, where *T* is the full set of cell types. We thus optimize *P_a_*(*t*) or 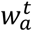 to minimize the total loss:

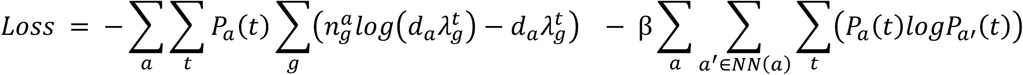

More details of the model and derivation of explicit form of gradient for fast optimization is available separately in **Supplementary File 1**. In short, we derived that the explicit form of gradient of the loss function ℒ at 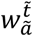 for spot *ã* and cell type *t̃* as:

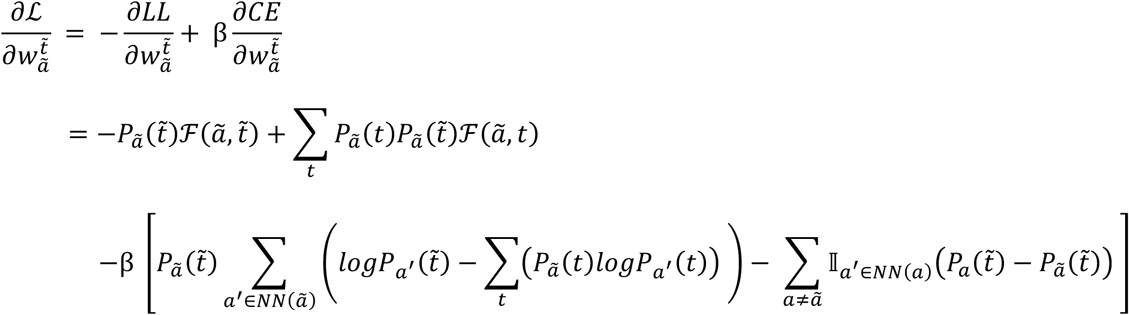

Where

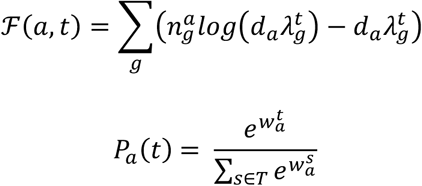

STHD thus optimizes 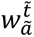 and the probability of spot *a*F having the identity of cell type 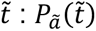, using Adam(16). We implemented iterative updates using numba(15) on sparse matrix to achieve high computational efficiency and scalable optimization.

### Construct normalized gene expression profile from reference single-cell atlas dataset

The average normalized gene expression 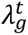 can be estimated from a pre-defined single-cell RNA-seq reference dataset. A reference single-cell dataset shall match the tissue or disease origin of the high resolution, whole-transcriptome spatial transcriptomics data. It shall also contain cell type annotation that is usually based on multi-sample harmonization and unsupervised clustering of single cells into various cell type or groups. The granularity of cell type annotation directly influence the number of predicted cell type and downstream analyses. For each cell type, average normalized gene expression profiles are estimated with 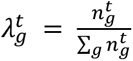 so that 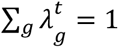. Specifically for human colon cancer, we downloaded the raw counts file of the single-cell RNA-seq data from the human colon cancer atlas from Pelka et.al(18) through GSE178341. The data contains 370115 tumor and adjacent normal single cells from 62 primary treatment-naïve colorectal cancer patients, with 43113 genes profiled. Quality control of single cells was performed by filtering cells with minimum 100 counts and 100 genes, filtering genes with minimum 3 cells, and removing cells with mitochondrial read ratio larger than 27.7%, which is mean plus one standard deviation of the mitochondrial read ratio across all cells. After the above quality control, 303358 cells and 31,873 genes remain. The cell annotations were obtained from the study that provided 98 fine-grained cell types and 7 cell lineages. We used the 98 high-granularity cell type annotation throughout this study. To select genes informative in distinguishing the 98 cell types, we followed the calculation of cell type expression profile as in RCTD(12). First, we calculated the estimated cell type expression profiles 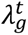 for all genes and cell types. We then selected genes with average expression profile above 0.000125, which is 0.0625 counts per 500, and at least 0.5 log-fold-change compared to the average expression across all cell types, using log2 scale. Moreover, we removed the mitochondrial genes and ribosome genes in the selection. The selection resulted in 4618 genes that are cell type-informative, for each we calculated 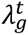 in all cell types. Note that STHD uses the 4618 genes to infer cell type identity of spots, and later can consider whole-transcriptome for downstream analyses.

### STHD hyperparameter tuning

To select hyperparameters that balance between statistical likelihood and neighborhood similarity, we comprehensively tested hyperparameter tuning and provided an automatic stopping criterion and optional tuning tutorial. The main parameters in STHD includes the neighborhood parameter β, which controls the contribution of cross entropy loss among local neighbors to the total loss, the optimization steps n_iteration, and the learning rate that can be fixed in practice. We provided a tutorial on parameter tuning based on performance in random sampled patches; for the human colon cancer sample, the optimized parameters are 23 iteration steps and β=0.1. In detail, for the colon cancer sample, we randomly selected six different patches in size of 1100×1100 pixels (in full resolution, around 22500 spots). For the optimization of each patch, we tested a range of β from 0, 0.03, 0.1, 0.3, 1, and 3, each outputting optimal iteration steps based on the early stopping criterion, where the drop of loss in the past 10 steps falls within 1%. Since the regularization term from neighborhood contribution β will lower log likelihood, as β increases, we select the optimal parameter pairs where drop in cross entropy slows down to around 10% while a high log likelihood is maintained. The optimal iteration steps when fixing β of 0, 0.03, 0.1, 0.3, 1, 3, is each 24, 23, 22, 23, 23, 24. The optimal iteration steps is robustly 23.17 on average. Therefore, we chose β=0.1 and n_iteration=23 as optimal parameters for the whole sample modeling that balance the count-based likelihood and the contribution from neighbors.

### STHD low-count region detection and filtering

Throughout the spatial transcriptomics data, regions of low gene counts exist, usually arising from tissue boundaries, tissue gaps, and regions low in DNA/RNA such as dead cells, necrosis, vessel enriching red blood cells, and collagen rich area. We included an automatic local low-count region detection and filtering functionality in STHD. By default, STHD computed neighbor graphs within 2 rings of spots (13 spots for the grid system), aggregated local connected spots, and detected low-count local regions not exceeding 50 counts, where the model will skip modeling due to low confidence in cell type inference. Other customized tissue gap and boundary masks can also be provided to further filter spots.

### Assessing model accuracy with simulated spatial transcriptomics data

To assess the accuracy of STHD model prediction, we simulated the high-dimensional, high-resolution spatial transcriptomics data *in silico*. We simulated a region with 22500 (150×150) grid spots and position 500 simulated cells overlapping the region with cell sizes spanning 1-20 spots. Each cell is randomly assigned a cell type from the 98 cell types annotated in colon cancer atlas, and spots overlapping each whole cell are assigned the corresponding cell type. To mimic the real-world scenario in gene expression profiles and spot sparsity, we adapted the same normalized gene expression profile from the same colon cancer atlas data with 98 cell types. The depth of each spot was simulated following the distribution of exponential decay with mean read depth 47 matching the colon cancer VisiumHD sample. After simulating counts of each gene at each spot in the region, we applied STHD with default optimal parameters (β=0.1) and also STHD no-neighbor model (β=0) to the simulated spatial data, and measured prediction accuracy, standard error, and area under the receiver operating characteristic curve (ROC), for this task of multi-class classification of 98 categories at each spot. To compared with other methods that enable spot-level cell type annotation, we examined six different settings: STHD, RCTD with default count filter, RCTD with no count filter, CARD with default count filter, CARD with no count filter, and Cell2Location. The inclusion of both default parameter and adjusted count filtering for RCTD and CARD is due to the fact that most spots will be removed if following default parameters, where accuracy measurements were based on non-filtered spots. We evaluated the methods in ten simulation replicates and assessed performance metrics in mean accuracy, AUC, and percentage of retained spots.

### Comparison with other spot cell typing methods

To benchmark the STHD results at 2×2 µm resolution, we compared STHD to six methods that can output cell typing, including the deconvolution-based cell typing tools RCTD(12), CARD(13), Cell2Location(14), Celloscope(20), Stereoscope(21), and the unsupervised method Banksy(22). Because of computational efficiency and data size limit of the existing methods, we used the colon cancer tumor region as in Fig.2a to perform benchmarking, which contains 22,336 cells and 18,085 genes. For methods requiring the reference scRNA-seq data as input, we used the same human colorectal cancer scRNA-seq datasets (GSE178341) with 98 cell types. Since many methods in comparison requires a maximum number of cells from reference scRNAseq dataset – for example, CARD requires below 50000 cells - we down-sampled the reference data to include maximum 600 cells for each cell type, resulting in a total of 49,351 cells and 41,195 genes. For running RCTD on single spots, we used RCTD doublet mode, testing both the default minimum UMI 100 and also by removing the minimum UMI requirement due to loss of most raw spots. For testing CARD, we followed guidelines from https://github.com/YMa-lab/CARD. Similarly, to avoid most spots being filtered, the minimum spot count and minimum gene numbers were set to 0. Cell2location and Stereoscope were ran with default parameters according to the tutorials https://cell2location.readthedocs.io/en/latest/notebooks/cell2location_tutorial.html, and https://github.com/almaan/stereoscope. In Cell2location, by default, the reference data was filtered using the following parameters: cell_count_cutoff = 5, cell_percentage_cutoff = 0.03, and nonz_mean_cutoff = 1.12. The reference single-cell regression model was trained with default parameter max_epochs = 250, lr = 0.002. The spatial cell2location model was obtained with default parameters max_epoch = 30,000. In Stereoscope, by default, the reference data was filtered using a minimum count threshold of min_counts = 10, and all mitochondrial genes were removed. The reference single-cell model was trained with parameters max_epochs = 100; the spatial stereoscope model was obtained using parameters max_epochs = 10,000. Finally, for Celloscope, a binary matrix with prior knowledge about marker genes is required among input, for which we selected top 200 genes in normalized expression from our informative gene matrix for each cell type. Another required input is estimated number of cells in each spot, for which we set to 1 due to sub-cellular-resolution nature of the data. Celloscope was then tested following guidelines at https://github.com/szczurek-lab/Celloscope. Finally, Banksy was tested with default parameters following the tutorial at https://github.com/prabhakarlab/Banksy_py/blob/main/slideseqv2_analysis.ipynb. For all toolkits, the weights or cell type abundance for each spot were extracted and normalized as cell proportions for subsequent visualizations.

### Cell type deconvolution in aggregated bins

Following current standards of binning-deconvolution, we binned the spatial data and aggregated the bins containing 4×4 spots. We applied RCTD deconvolution of doublet mode on the same spatial patch as in Fig.2a, following the default filtering and parameters of RCTD as on spacexr (https://github.com/dmcable/spacexr), and using the same subsampled colon cancer reference scRNA-seq data as in comparison with other methods. To compare cell proportions across bins, we counted STHD cell type labels and obtained cell proportions for each 4×4 bin after excluding filtered and ambiguous spots. We also computed cosine similarity between STHD’s spot count-based proportions and RCTD’s deconvoluted proportions and visualize the proportions and cosine similarity on a spatial plot.

### Estimating running time in computational efficiency

To assess computational efficiencies, different cell typing tools are applied onto the tumor epithelial patch using a server with 8 CPU cores and maximum 128GB memory, running time recorded. For tools supporting GPU acceleration (cell2location and stereoscope) the time cost was evaluated on a single GPU, and the spatial training time was recorded without including scRNA training time. For computational efficacy, we also estimated the time cost for the entire sample by linearly scaling running time based on number of spots in patch (22,336 spots) and in the entire colon cancer sample P2 (8,726,224 spots).

### STHD pipeline on an entire VisiumHD sample

The STHD software and pipeline contains several components to flexibly split the whole sample into patches for inferences and parallelization. The input/output functionalities (STHDio) expanded from Squidpy Visium interface to enable cropping of full-resolution image. Second, for the full-size sample, STHD will automatically patchify the full sample into multiple regions with overlaps, followed by low-count filtering, per-patch inference, and result aggregation where probabilities of overlapping spots are averaged. The STHD cell type label results are determined by Maximum A Posterior (MAP) with a preset cutoff (default 0.8). An optional fast implementation for STHD-guided binning with a predefined bin size is provided, which outputs per-gene count for each bin stratified by STHD cell type labels. Finally, the predicted cell labels are exported to STHDviewer for a rasterized and interactive scatter plot implemented using Bokeh(49). In STHDviewer, the spot grid of array_row and array_column were used for x-axis and y-axis for visualization efficiency.

### Cell type-specific differential gene expression analysis across spatial regions of interest

To study regional differences within the cancer sample, we first focused on regions enriching epithelial cell types and manually separated four different regions and extracted spot barcodes. Based on STHD predicted labels, we performed differential expression analyses specific for epithelial cell types. Gene counts were normalized towards the spot count depth (target total counts 10000) and log-transformed followed by Wilcoxon test in Scanpy. Genes passing adjusted p-value cutoff 10^34^and absolute log2 fold change 0.5 were selected as significant, and top genes passing log2 fold change 0.5 are exported to pathway enrichment analyses by GSEApy(50) using EnrichR and a few pathway databases, including GO Molecular Function 2023, GO Biological Process 2023, MSigDB Hallmark 2020, Reactome 2022, WikiPathway 2023 Human, KEGG 2021 Human, and BioPlanet 2019. The FDR cutoff of 0.1 was used to filter the pathways and top four pathways were visualized for tumor epithelial pathway comparison.

We also tested cell type-specific differential expression using both STHD-labeled spot and STHD-guided bin aggregates of size 4×4 spots. At STHD-guided bin level, we followed the same step of normalization and performed differential expression with Wilcoxon with the same cutoff for epithelial and macrophage comparison. For T cell-specific differential expression analyses between the two T cell rich regions, the cutoff was set log2 fold change 0.5 and adjusted p-value cutoff 0.05 due to small number of spots from patches. For differential gene dotplots, significant genes are sorted by log fold change and the top three genes for the tumor comparison and top ten genes for the immune comparison were plotted.

### Neighborhood interaction enrichment analysis for immune-rich regions in colon cancer VisiumHD sample

To investigate the neighborhood interaction patterns across various cell types in the two immune-infiltrated regions, local neighborhoods were calculated using the 4 nearest cells in the grid and considering 6 rings of adjacent spots to get a obtain a local neighborhood structure using squidpy.gr.spatial_neighbors in Squidpy. We used STHD predicted cell type labels and excluded spots in the filtered and ambiguous categories. The neighborhood enrichment Z-scores were calculated using squidpy.gr.nhood_enrichment, which follows CellChat(38) with ligand-receptor databases and permutation of 1000 times. Heatmaps of neighborhood interaction patterns were plotted based on the co-occurrence score of pairwise cell types in each region of interest.

### Inter-cell type frontline and cell-cell communication analysis

For a given pair of cell types, we first defined their precise frontlines to be spots where those two cell types are adjacent to each other. We further restrict frontline spots to be those whose neighbor only contains that pair of cell types based on maximum posterior to eliminate potential noises from other cell types. Considering a single cell is comparable to the size of 4 spots, we assumed spots within 4 spots from the frontlines represented one cell with direct interaction with another cell type. Thus, spots were categorized based on their cell type and proximity to frontline. In the colon cancer sample P2, we examined interactions between T cells and dendritic cells, and T cells and macrophages separately for the two immune regions of interest, Immune-Trich-left and Immune-Trich-right. In the chord plots, for a pair of cell types A and B, the labels were assigned as follows: ‘1’ for far-frontline cell type A, ‘2’ for far-frontline cell type B, ‘3’ for near-frontline cell type A, and ‘4’ for near-frontline cell type B. We then hypothesized that cell-cell communications like ligand-receptor expression levels were higher between group 3 and 4 compared to group 1 and 2. To perform the ligand-receptor interaction analysis, normalized data was applied the squidpy.gr.ligrec function which permuted 1000 times to detect significant ligand-receptor pairs. Top 10 ligand-receptor pairs different between near-frontline and far-frontline communications were visualized in the chord diagram showing the mean expression of each ligand-receptor pair and sender-receiver directions.

### STHD on additional colon samples and other human and mouse tissue samples

We performed STHD analyses on the extended list of human colon samples using the same colon cancer reference gene estimation, yielding cell type inference on 8,115,201 spots in colon cancer P1, 8,665,510 spots in colon cancer P5 sample, and 7,791,374 spots in the normal colon sample P3 using normal cell types. We further extended analyses to the human pancreas sample containing 6,680,357 spots for 18,085 genes, the mouse intestine sample containing 5,479,447 spots with 19,059 genes, and the mouse brain sample containing 6,296,637 spots with 19,059 genes. All analyses were run using default parameters. To match the tissue system, we used the non-cancerous colon scRNA for the normal colon sample, normal human adult pancreas scRNA dataset for the human pancreas sample, the adult Mouse Cell Atlas intestine scRNA dataset for the mouse small intestine sample, and the Allen Brain Atlas reference for the mouse brain sample.

For constructing reference gene expression profile for human pancreas, we downloaded raw counts file of single-cell RNA-seq data from the human pancreas atlas by Baron et al.(41) from GSE84133. The dataset contains 8,569 normal single cells from four donors, profiling 20,124 genes. Quality control involved filtering cells with a minimum of 100 counts and 100 genes, filtering genes with a minimum of 3 cells, and removing cells with a mitochondrial read ratio larger than 5%. After quality control, 8,569 cells and 16,358 genes remained. The study provided 14 cell type annotations. Using STHD default setting, we selected 4,436 cell type-informative genes based on average expression profile above 0.000125 (0.0625 counts per 500) and at least 0.5 log-fold change, excluding mitochondrial and ribosomal genes. For mouse intestine tissue, the raw counts file of single-cell RNA-seq data from the mouse cell atlas by Fei et al.(42) was downloaded from GSE176063. The dataset contains 6,684 normal single cells from three adult mouse intestine samples, profiling 16,416 genes. Quality control involved filtering cells with a minimum of 100 counts and 100 genes, filtering genes with a minimum of 3 cells, and removing cells with a mitochondrial read ratio larger than 7.15%. After quality control, 6,102 cells and 15,745 genes remained, with 28 cell types annotated. Similarly, we selected 4,083 cell type-informative genes based on average expression profile above 0.0002 (0.1 counts per 500) and at least 0.5 log-fold change, excluding mitochondrial and ribosomal genes. For mouse brain reference gene expression construction, the processed single-cell RNA-seq data from the Allen Brain Atlas was downloaded from the Seurat tutorial website (https://satijalab.org/seurat/articles/visiumhd_analysis_vignette), which contained 199,993 single cells and 31,053 genes. Quality control involved filtering cells with a minimum of 100 counts and 100 genes, filtering genes with a minimum of 3 cells, and removing cells with a mitochondrial read ratio larger than 4.8%. After quality control, 181,250 cells and 264,484 genes remained, with 41 subclass cell types annotated. Similarly, we selected 5,171 cell type-informative genes based on average expression profile above 0.000125 (0.0625 counts per 500) and at least 0.5 log-fold change, excluding mitochondrial and ribosomal genes.

## Declarations

### Availability of data and materials

#### Software availability

The current STHD software source code is on GitHub at https://github.com/yi-zhang/STHD.

An interactive STHDviewer of the high-resolution cell type result for the human colon cancer sample P2 is available at https://yi-zhang-compbio-lab.github.io/STHDviewer_colon_cancer_hd/STHDviewer_crchd.html.

#### Data Availability

All analyses in this paper are based on publicly available data.

Colon cancer reference scRNA-seq atlas: data and cell type annotation from Human Colon Cancer Atlas is available from Pelka et. al through GSE178341: (https://www.ncbi.nlm.nih.gov/geo/query/acc.cgi?acc=GSE178341).

Colon cancer VisiumHD data (patient P2) is available at https://www.10xgenomics.com/datasets/visium-hd-cytassist-gene-expression-libraries-of-human-crc, named Visium HD Spatial Gene Expression Library, Human Colorectal Cancer (FFPE), by Space Ranger 3.0.0, 10x Genomics (2024, Mar 25).

Colon cancer VisiumHD data (patient P1, P3, P5) from Oliveira et al.(9) include two colon cancer and one normal colon, available at https://www.10xgenomics.com/products/visium-hd-spatial-gene-expression/dataset-human-crc.

The human pancreas VisiumHD data is available at https://www.10xgenomics.com/datasets/visium-hd-cytassist-gene-expression-libraries-human-pancreas.

The mouse small intestine VisiumHD data is available at https://www.10xgenomics.com/datasets/visium-hd-cytassist-gene-expression-libraries-of-mouse-intestine.

The mouse brain VisiumHD data is available at https://www.10xgenomics.com/datasets/visium-hd-cytassist-gene-expression-libraries-of-mouse-brain-he.

### Authors’ contributions

This study was conceived and led by Y.Z. Y.Z. designed the model and algorithm, implemented the STHD software, led the data analyses, and wrote the manuscript. C.S. analyzed the data and helped with manuscript writing.

## Acknowledgements

We acknowledge 10X Genomics to share VisiumHD datasets publicly available. We acknowledge authors of the Human Colon Cancer Atlas to make scRNAseq datasets publicly available.

