## Supplementary Figures for "STHD: probabilistic cell typing of single Spots in whole Transcriptome spatial data with High Definition"

**Supplementary Fig.1.**

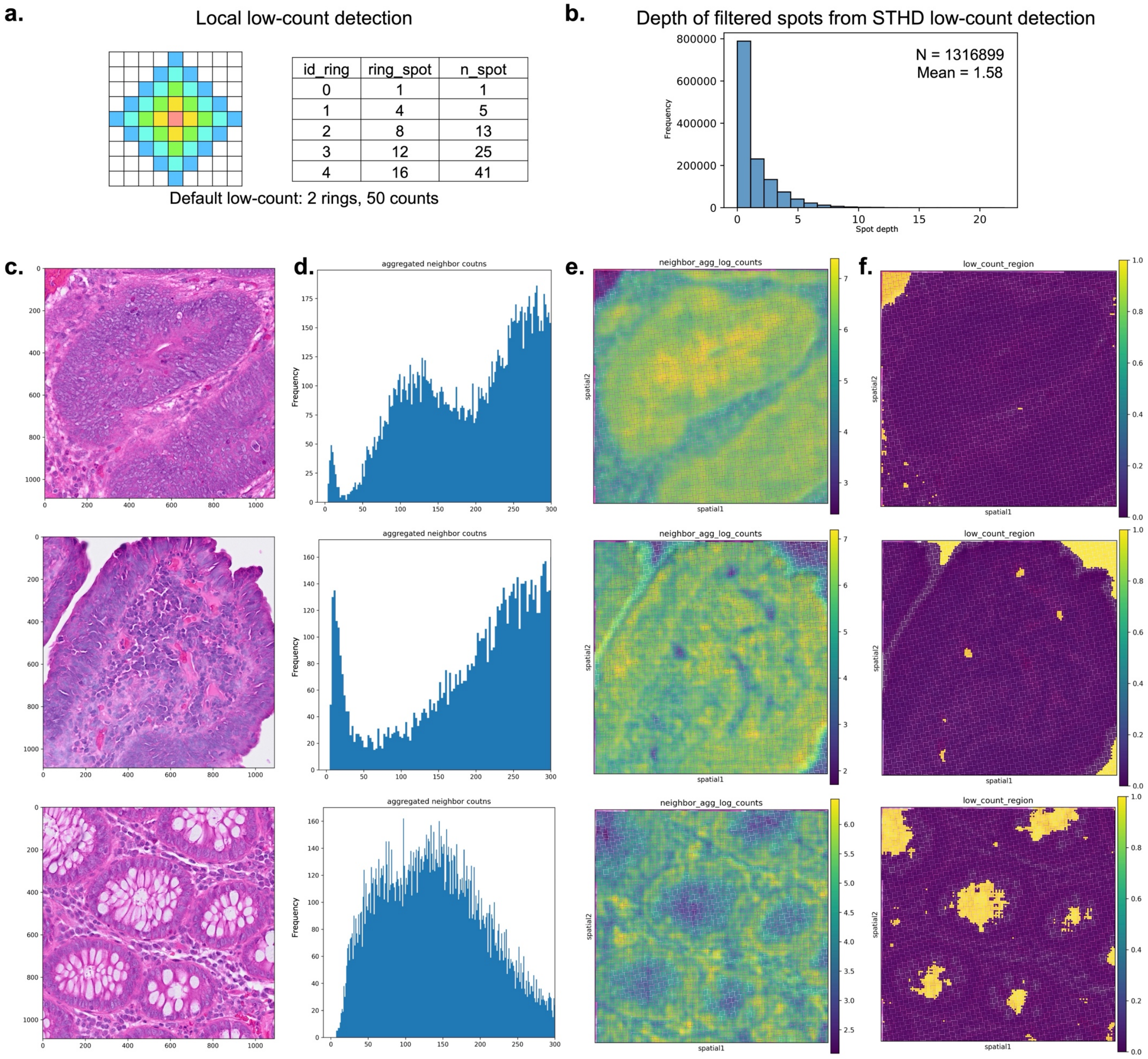


**Supplementary Fig.1. Low-count detection in STHD. a.** Illustration of the number of neighboring spots considered in local low-count detection when varying ring numbers. **b.** The depth distribution of spots detected as low-count region for the colon cancer VisiumHD sample P2. **c.** H&E histology images showcasing three example regions in the colon cancer VisiumHD sample P2 with different morphology. **d.** The corresponding distributions of local aggregated spot counts within each patch. **e.** Heatmap of neighbor-aggregated counts showing locally continuous low-count regions in the corresponding patch. **f.** The detected spots assigned as low-count regions, which are later filtered in STHD using default thresholding parameters.

**Supplementary Fig.2.**

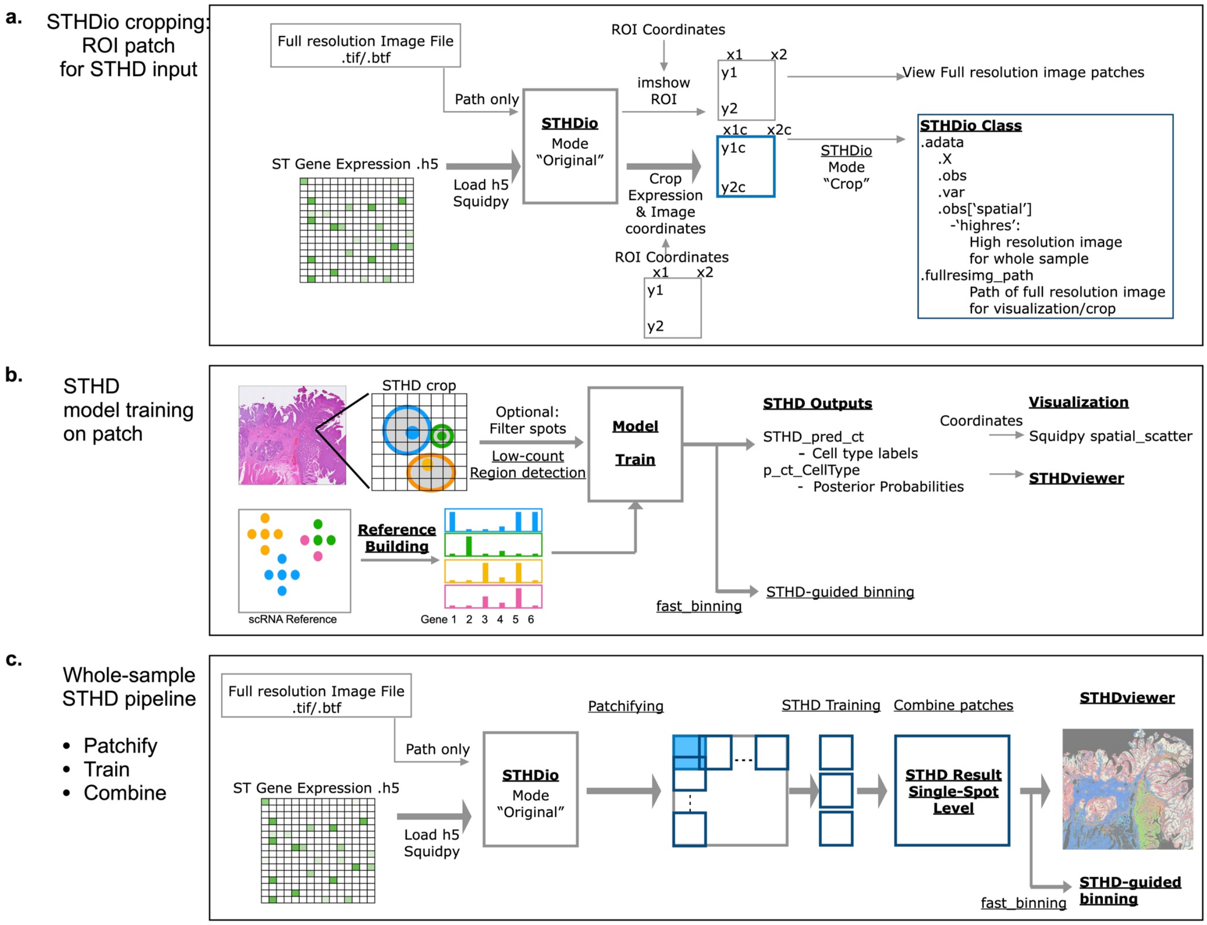


**Supplementary Fig.2. STHD implementation. a.** The Squidpy-based input/output design in STHD, for efficient and flexible cropping of spatial sample based on coordinates following full-resolution images. The input coordinates are adjusted to fit intact spot regions. **b.** STHD training steps at the patch level with input of pre-built gene expression profile from reference scRNA-seq data. The outputs are in format of cell type labels and per-cell type posterior probabilities at spot level. **c.** STHD pipeline applied to the entire sample with automatic patch parallelization. ROI, region of interest.

**Supplementary Fig.3.**

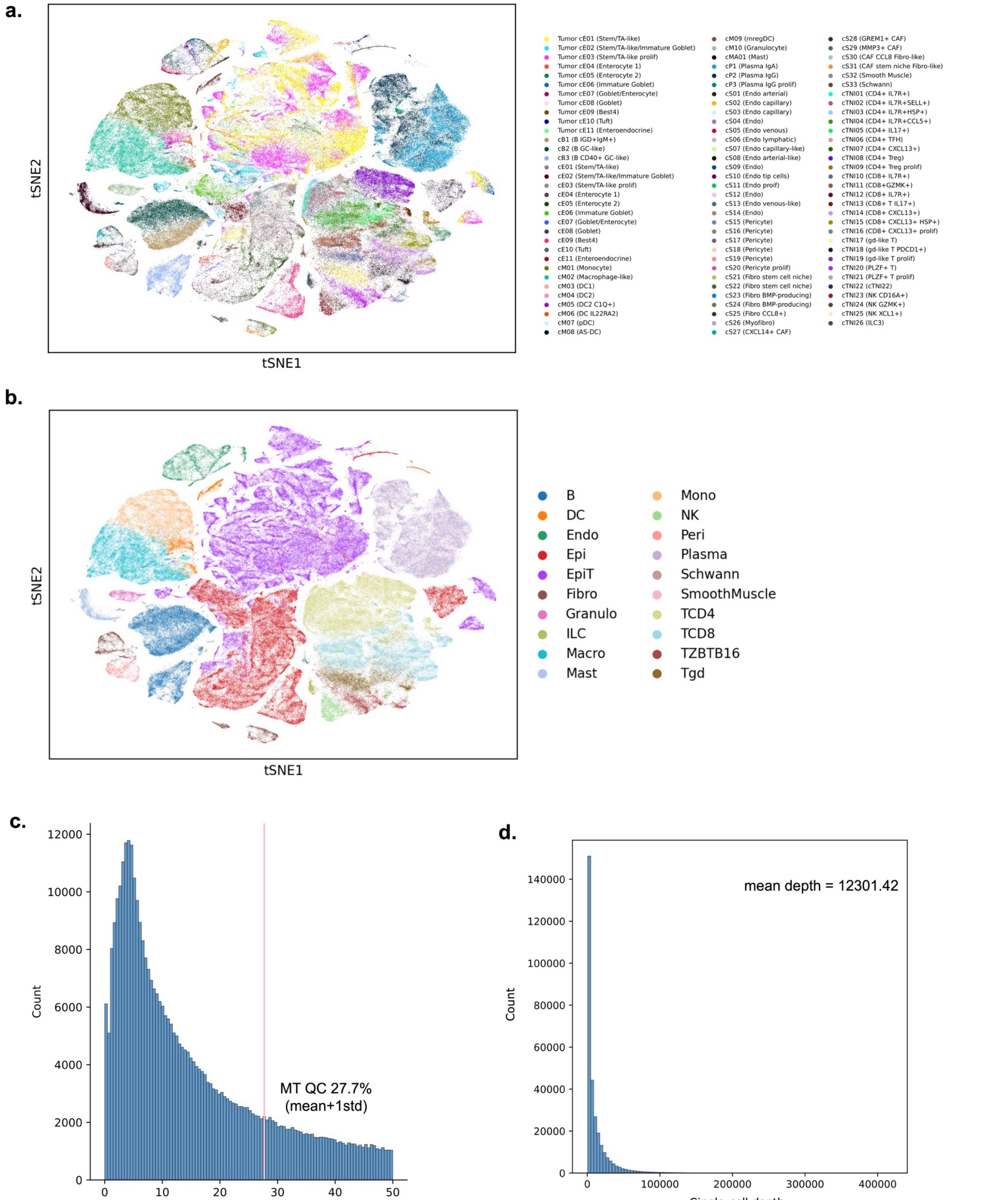


**Supplementary Fig.3. Colon cancer reference scRNA dataset. a.** The reference scRNA-seq atlas from human colon cancer atlas after QC on mitochondrial reads. Cells are colored by the 98 cell types annotated in the original study. **b.** Distribution of mitochondrial read ratios of the raw data. **c.** The count depth distribution of all single cells in the reference scRNA dataset.

**Supplementary Fig.4.**

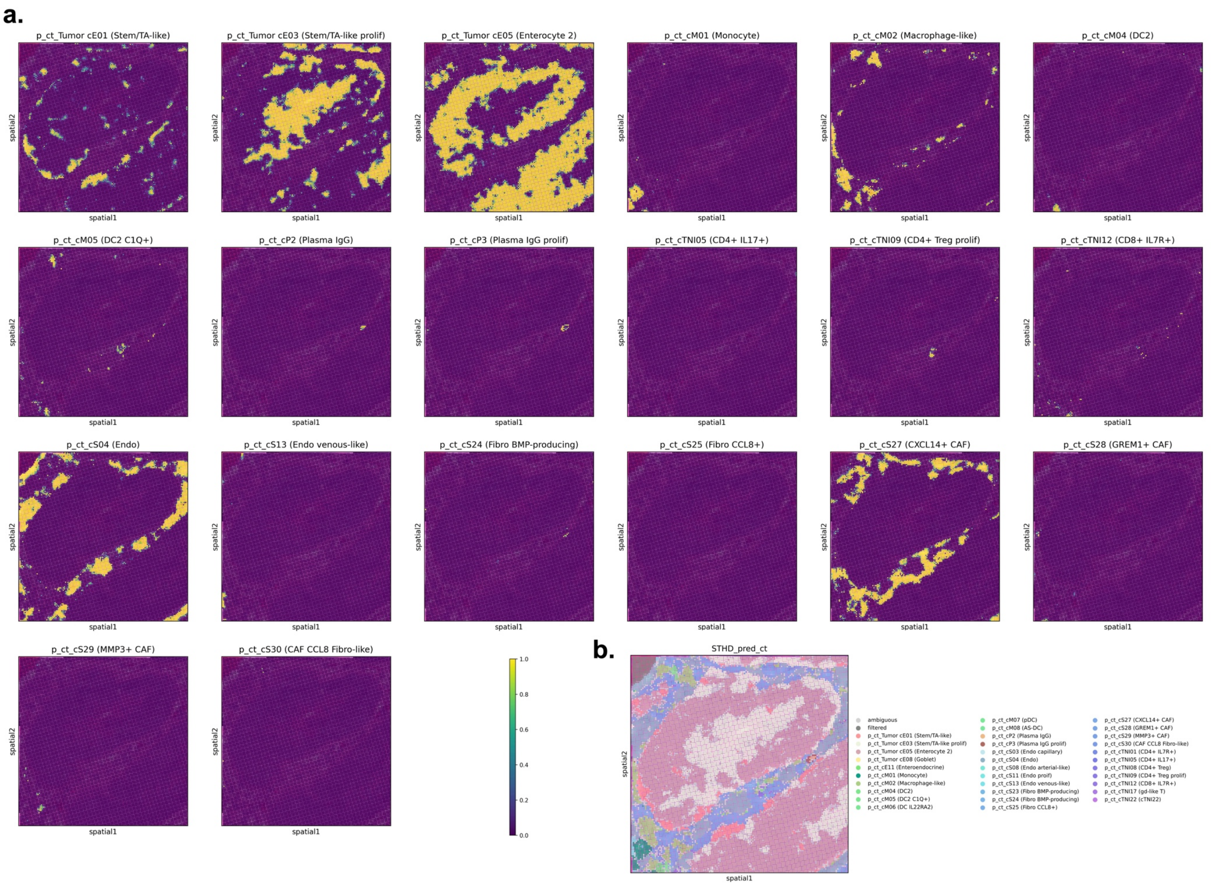


**Supplementary Fig.4. STHD cell type probabilities per spot. a.** STHD spot-level cell type probabilities for the tumor epithelial patch visualized using Squidpy. Cell types with at least 3 spots in patch are retained to show cell type posterior probabilities. **b.** STHD predicted cell type labels at spot level visualized with Squidpy.

**Supplementary Fig.5.**

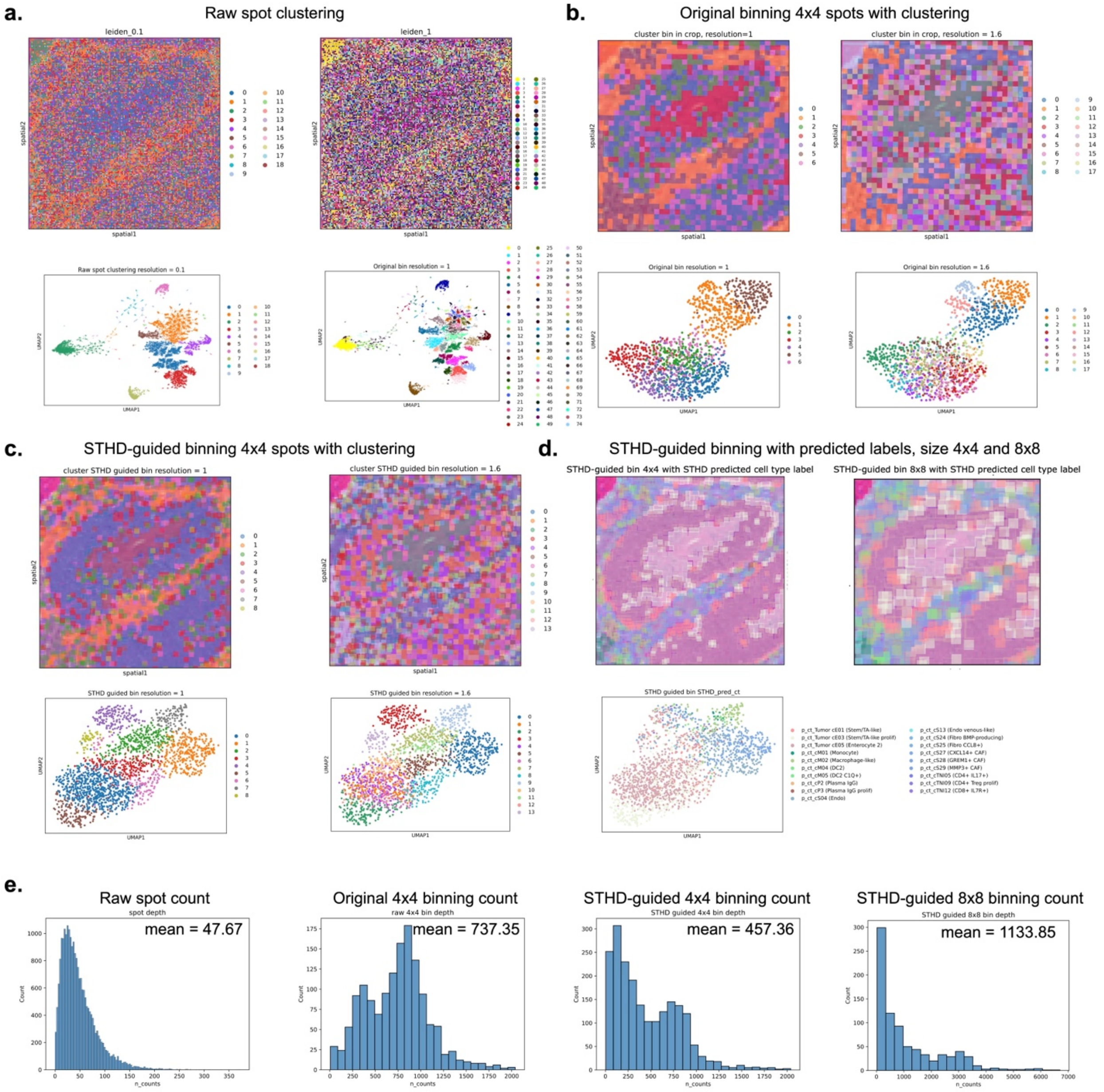


**Supplementary Fig.5. Showcase of different computational strategies for high-resolution cell typing on the test patch. a.** Spatial scatter plot and spot UMAP plot after clustering spots using different resolution. **b.** Clustering aggregated bin-level gene expression in size of 4x4 spots, using different clustering resolution. **c**. Clustering aggregated bins stratified by STHD cell type labels for further transcriptional analyses, using different clustering resolution. **d.** Visualizing STHD-guided bin using Squidpy spatial scatter plot, aggregating 4x4 spots and 8x8 spots per STHD-guided bins. **e**. Count depth distributions at spot-level, original bin level (4x4 spots), and STHD-guided bin levels (4x4 spots or 8x8 spots stratified by cell type), for the test patch.

**Supplementary Fig.6.**

**a.**

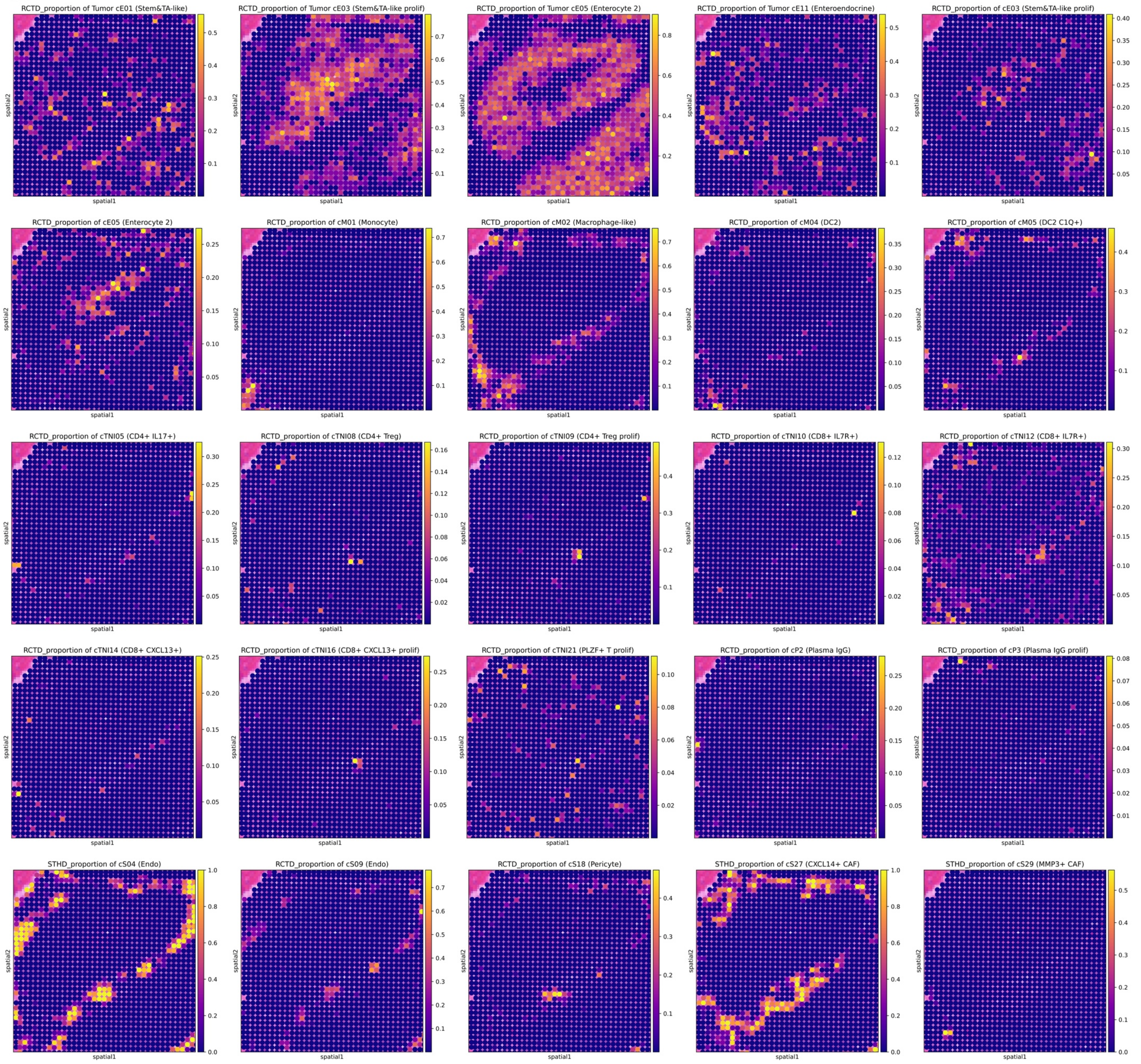


**b.**

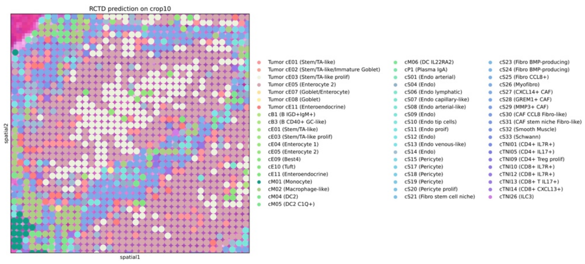


**c.**

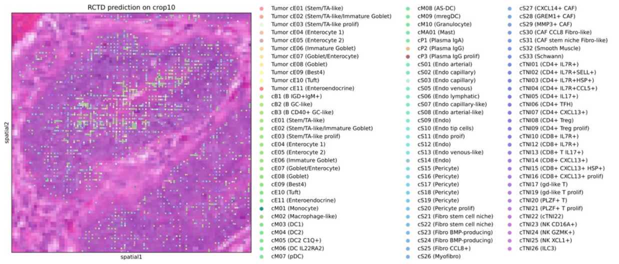


**Supplementary Fig.6. Testing deconvolution-based cell typing on bins and spots. a.** Cell type proportions by RCTD full-mode deconvolution on the 8x8um bins in size of 4x4 spots. **b.** Cell type labeling using rank-one cell type based on RCTD doublet mode on the 8x8um bins in size of 4x4 spots. **c.** Testing cell type labeling by RCTD doublet mode on 2x2um size spots with default UMI filtering.

**Supplementary Fig.7.**

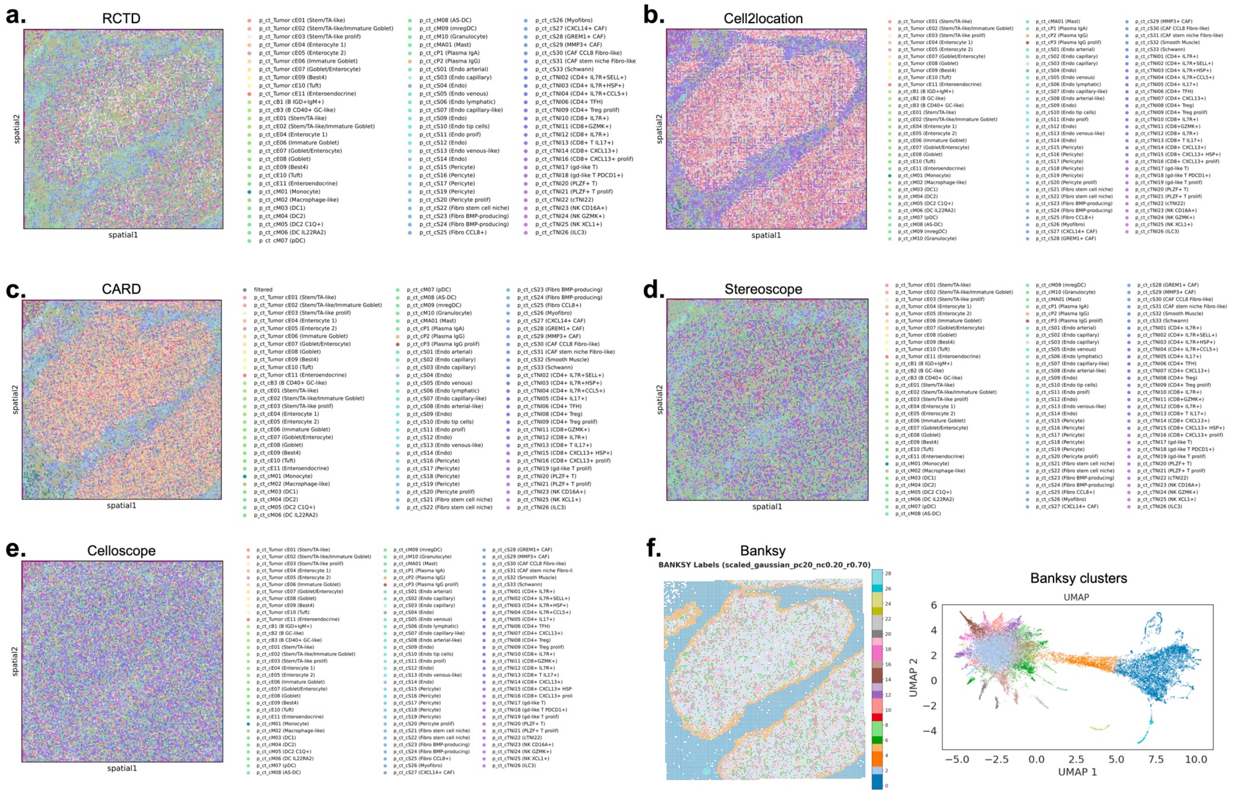


**Supplementary Fig.7. Comparison with available computational methods that generate cell type labels at spot level. a.** Rank-one cell type from RCTD doublet mode on spot-level expression profiles for the tumor epithelial patch. The default UMI filtering was removed to obtain results for all spots. **b.** Cell type labeling by Cell2location. **c.** Top cell type from CARD. **d.** Top cell types from Stereoscope. **e.** Cell type from Celloscope. **f.** Spot clustering using Banksy with spot UMAP embeddings.

**Supplementary Fig.8.**

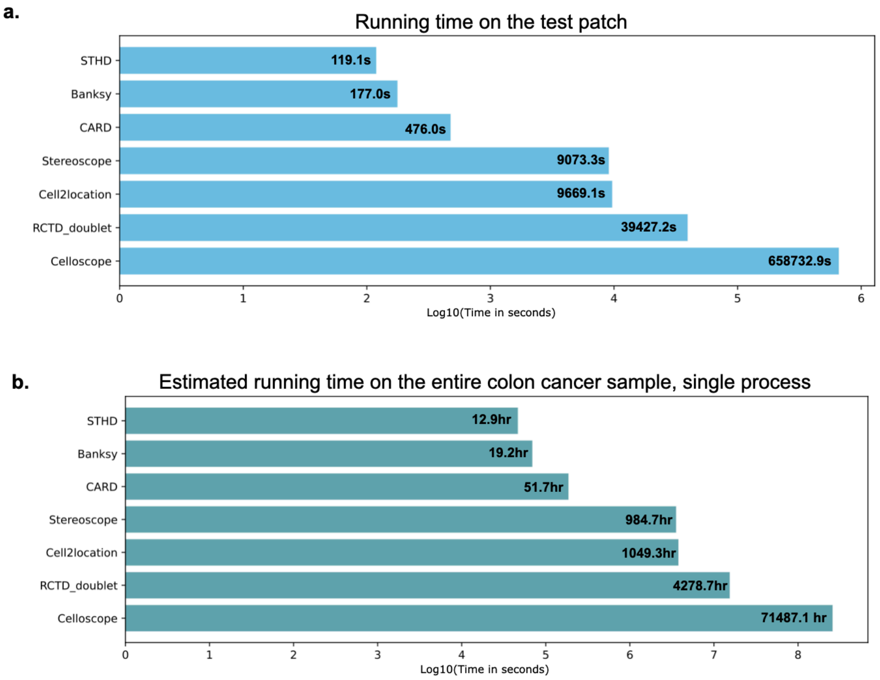


**Supplementary Fig.8. Running time. a.** Time cost for inferring cell type labels for 22336 spots from the 300x300um tumor epithelial test patch, ran on a single 8-core CPU. For Cell2location, the training time for reference single-cell RNA was not included. For Celloscope, the default number of sampling steps was followed. **b.** The estimated running time for the entire sample by linearly scaling number of spots in patch to the entire sample. The time estimation is based on running on a single 8-core CPU processor.

**Supplementary Fig.9.**

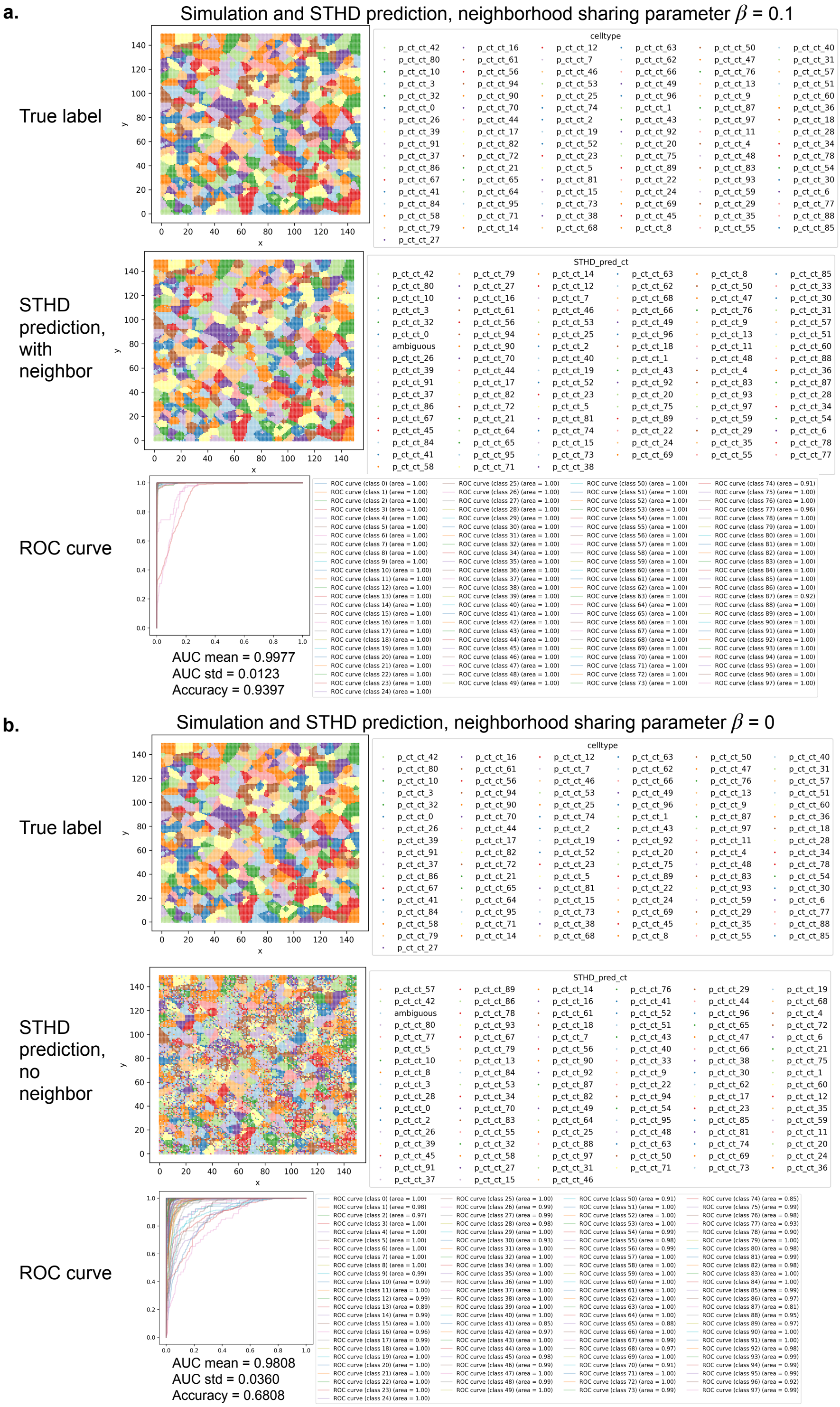


**Supplementary Fig.9. Benchmarking with simulated spatial data. a.** Top, simulated spatial data with each spot colored by true cell type label. Middle, STHD predicted cell type labels. Bottom, ROC curve of the multi-class classification. **B.** Top, simulated spatial data with each spot colored by true cell type label. Middle, STHD predicted cell type labels without contribution from neighbor spots. Bottom, ROC curve of the multi-class classification. ROC: receiver operating characteristic curve.

### Supplementary Fig.10.

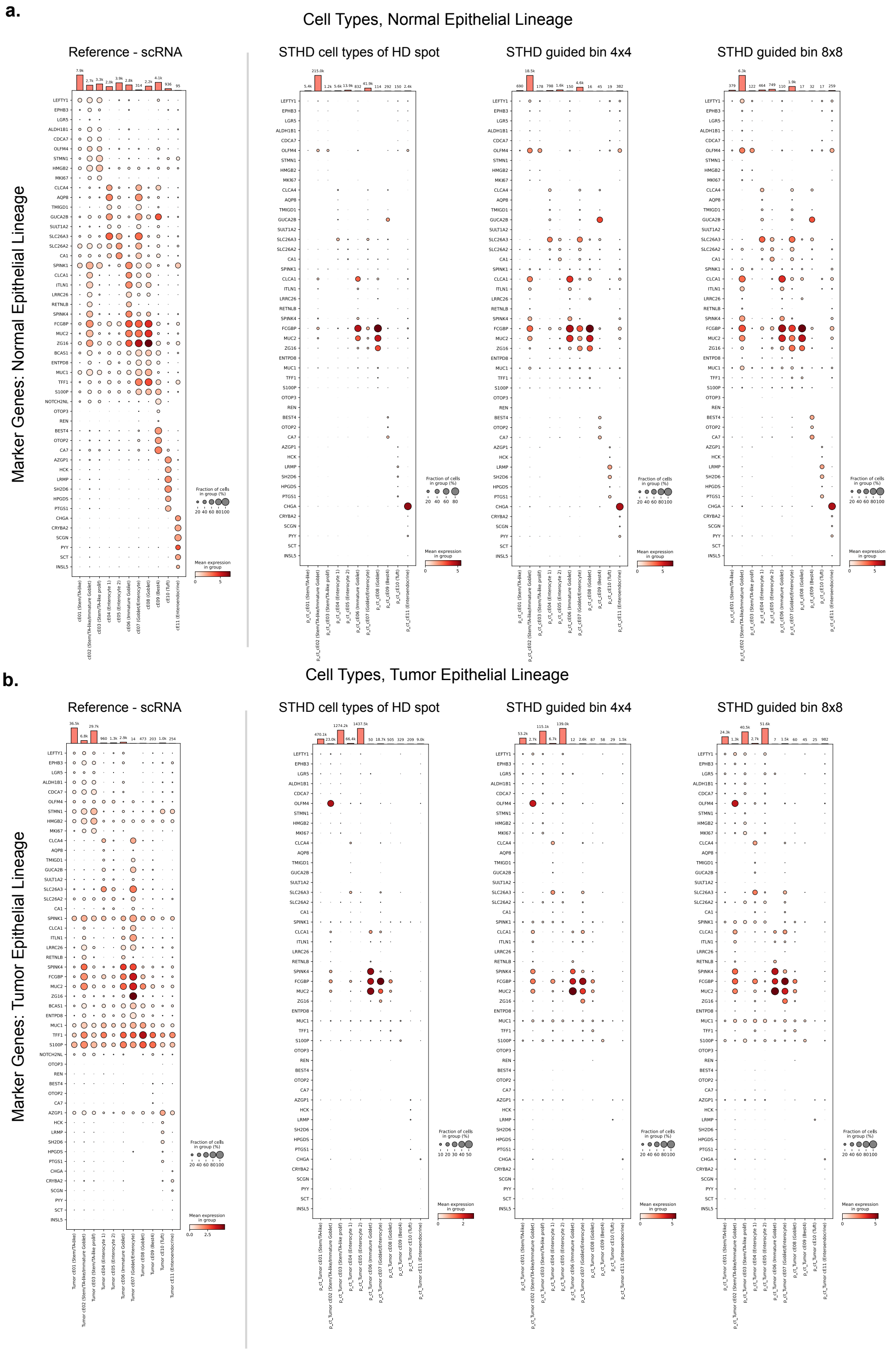


**Supplementary Fig.10. Expression dot plots for marker genes of normal and tumor epithelial cells.** Left to right: epithelial cell types in reference human colon cancer sample, epithelial cell types based on STHD predicted spots, epithelial cell types from STHD-guided bins of size 4x4 spots, epithelial cell types in STHD-guided bins of size 8x8 spots. **a.** normal epithelial markers and types; **b.** tumor epithelial markers and types. Marker genes and cell types are extract from the original single-cell study.

**Supplementary Fig.11.**

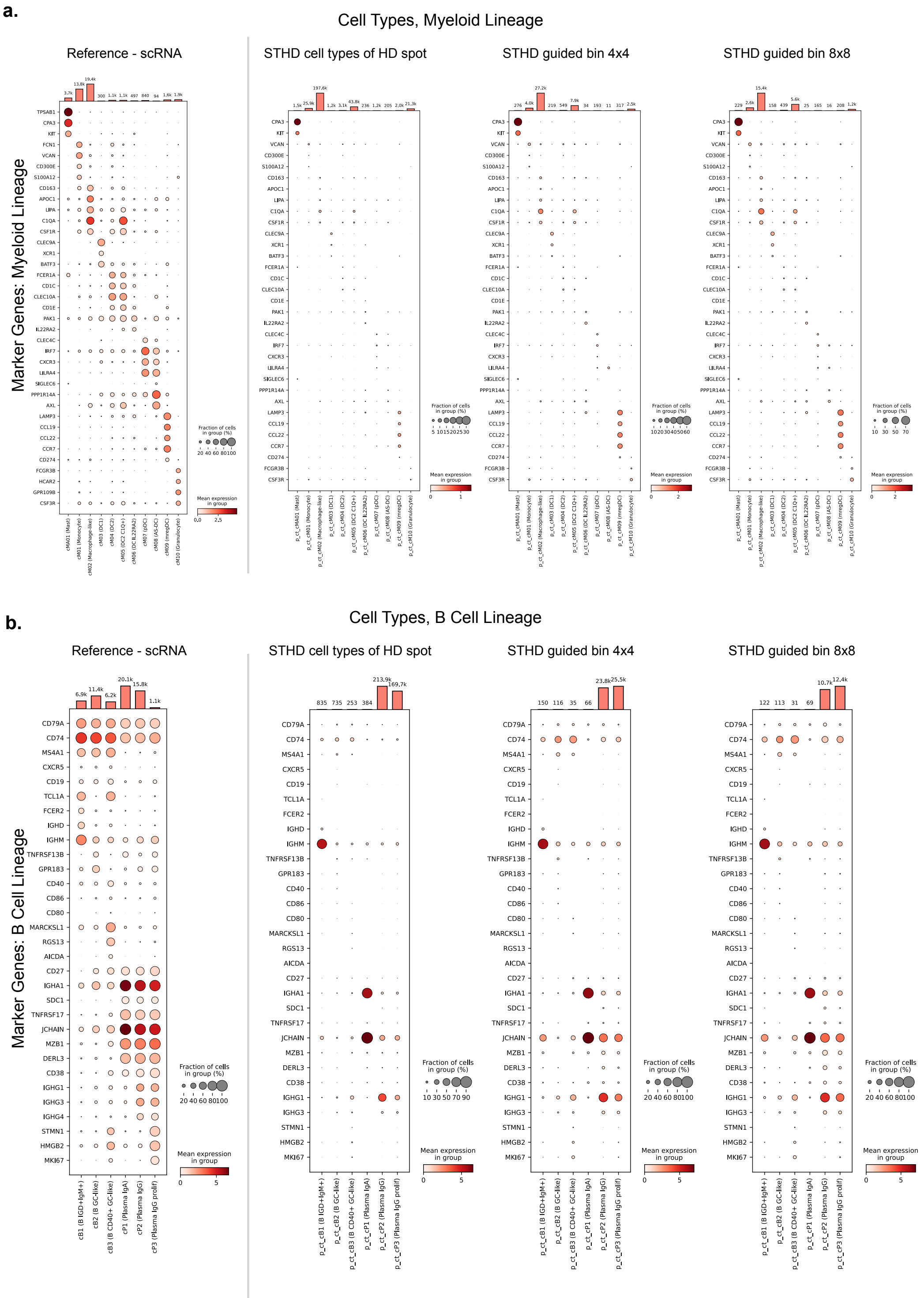


**Supplementary Fig.11. Expression dot plots for marker genes of myeloid and B cells.** Left to right: reference human colon cancer sample, cell types based on STHD predicted spots, cell types from STHD-guided bins of size 4x4 spots, cell types in STHD-guided bins of size 8x8 spots. **a.** myeloid markers and types; **b.** B cell markers and types. Marker genes and cell types are extract from the original single-cell study.

**Supplementary Fig.12.**

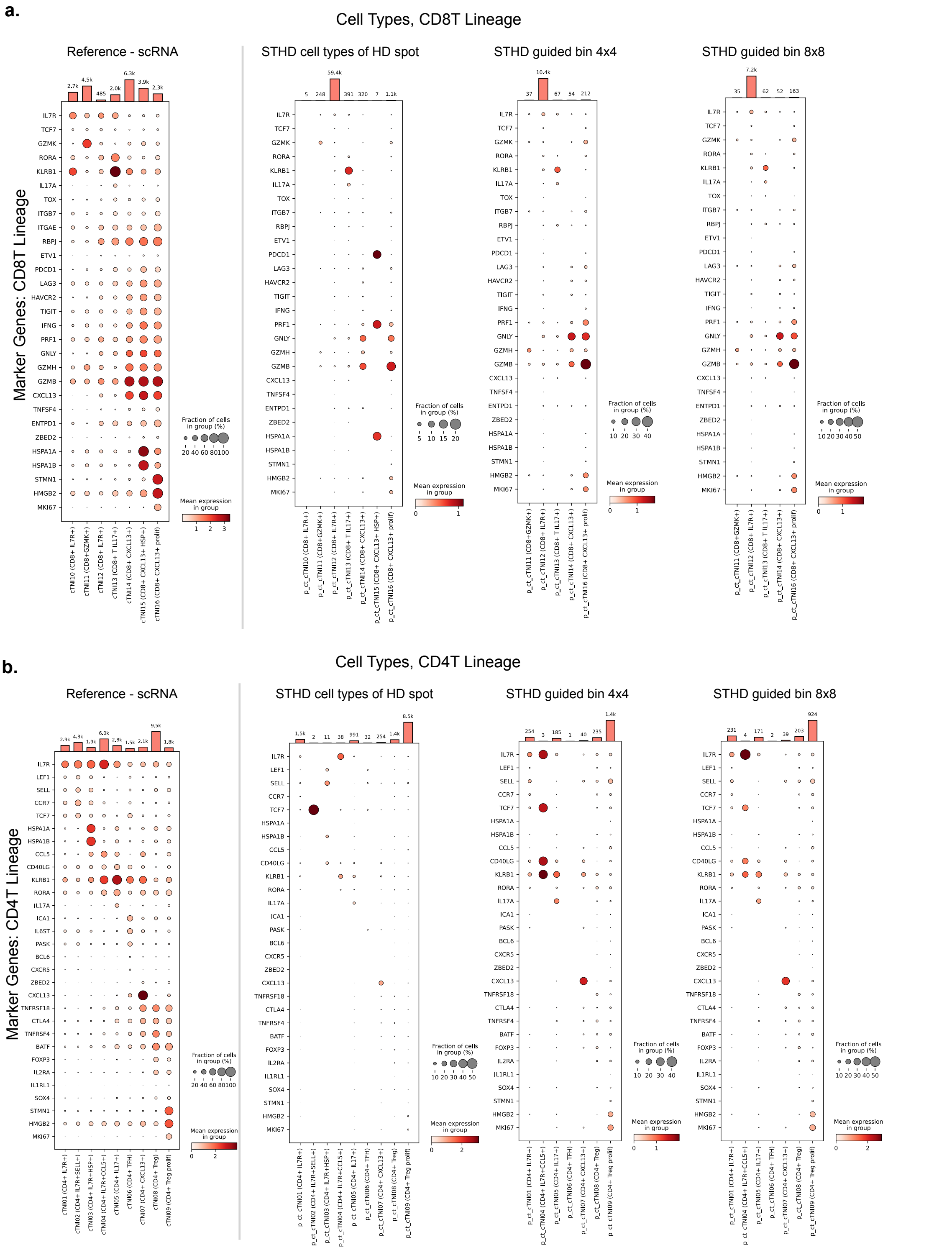


**Supplementary Fig.12. Expression dot plots for marker genes of T cells.** Left to right: reference human colon cancer sample, cell types based on STHD predicted spots, cell types from STHD-guided bins of size 4x4 spots, cell types in STHD-guided bins of size 8x8 spots. **a.** CD8T markers and types; **b.** CD4T cell markers and types. Marker genes and cell types are extract from the original single-cell study.

**Supplementary Fig.13.**

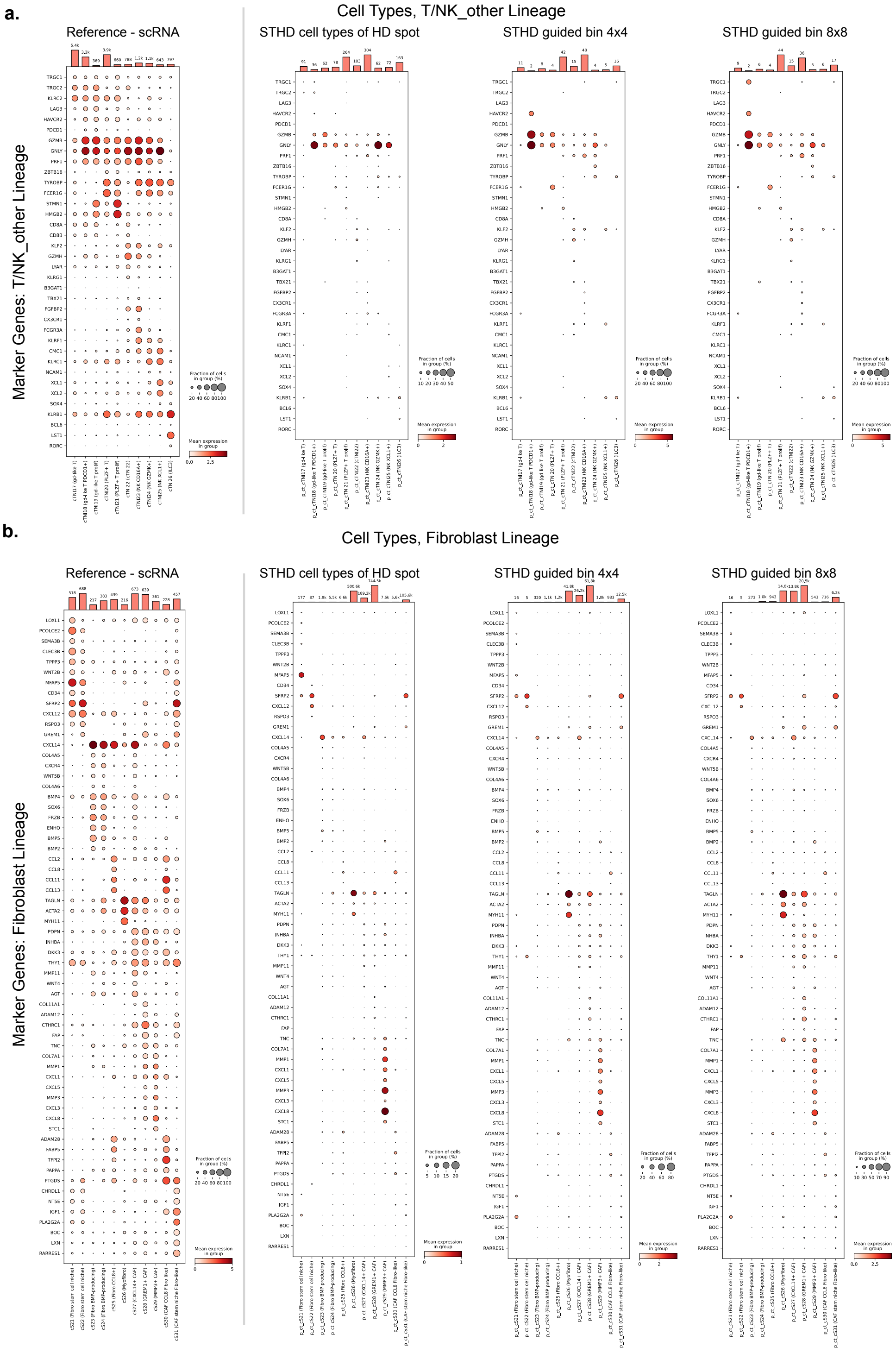


**Supplementary Fig.13. Expression dot plots for marker genes of other T/NK cells and fibroblast cells.** Left to right: reference human colon cancer sample, cell types based on STHD predicted spots, cell types from STHD-guided bins of size 4x4 spots, cell types in STHD-guided bins of size 8x8 spots. **a.** other T cell or NK cell markers and types; **b.** fibroblast cell markers and types. Marker genes and cell types are extract from the original single-cell study.

**Supplementary Fig.14.**

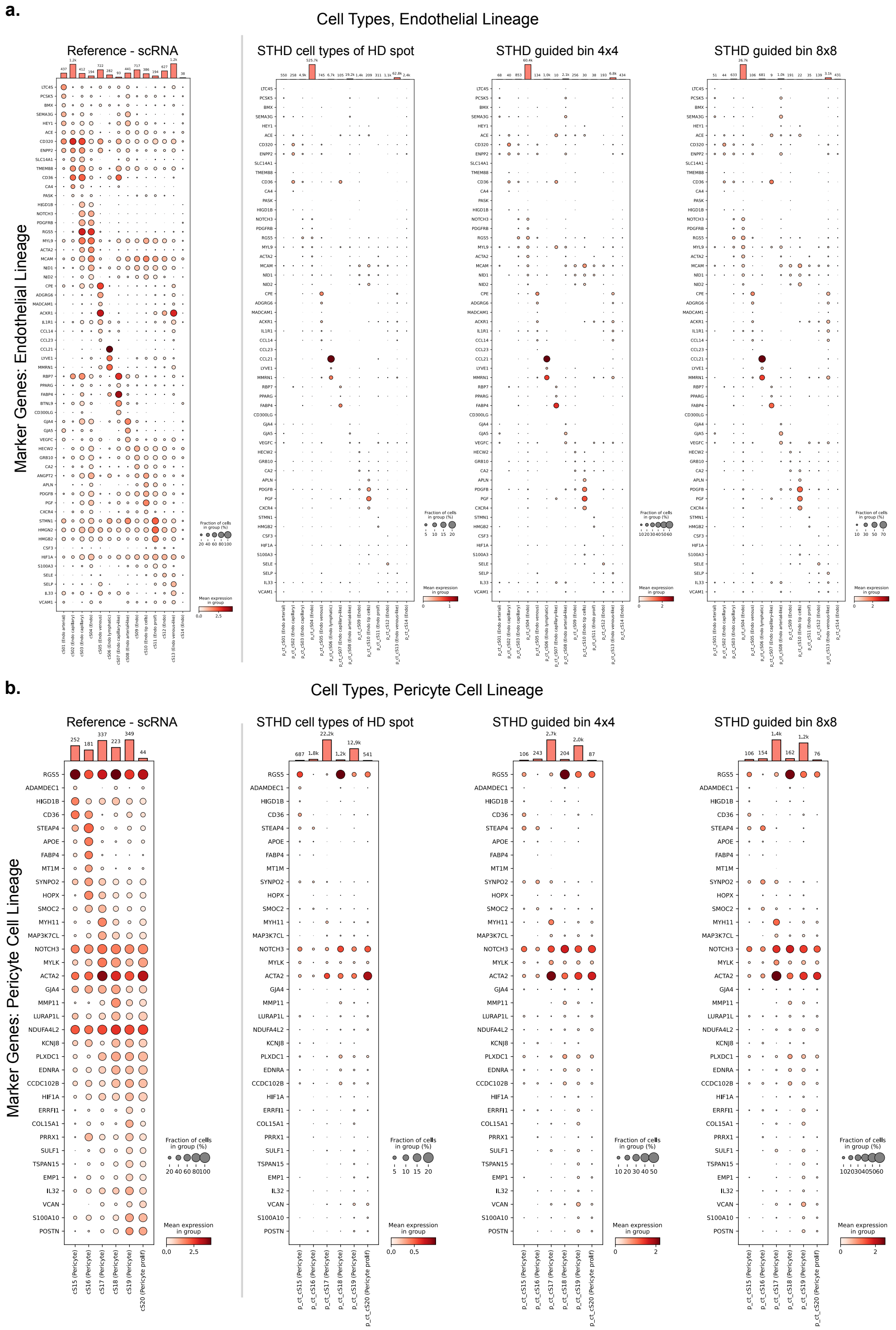


**Supplementary Fig.14. Expression dot plots for marker genes of endothelial and pericyte cells.** Left to right: reference human colon cancer sample, cell types based on STHD predicted spots, cell types from STHD-guided bins of size 4x4 spots, cell types in STHD-guided bins of size 8x8 spots. **a.** endothelial cell markers and types; **b.** pericyte cell markers and types. Marker genes and cell types are extract from the original single-cell study.

### Supplementary Fig.15.

**Patch 1**

Coordinates:
x1=52500
x2=53600
y1=9250
y2=10350

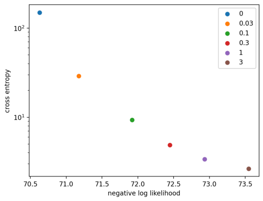


| beta | n_iter | LL | CE | ratio_CE_ lower_LL | CE_drop |
|---|---|---|---|---|---|
| 0 | 22 | -70.63 | 149.44 | 0.000 | 1.000 |
| 0.03 | 22 | -71.17 | 29.05 | 0.008 | 0.194 |
| **0.1** | **24** | **-71.92** | **9.36** | **0.018** | **0.063** |
| 0.3 | 27 | -72.45 | 4.88 | 0.026 | 0.033 |
| 1 | 33 | -72.93 | 3.39 | 0.033 | 0.023 |
| 3 | 54 | -73.55 | 2.65 | 0.041 | 0.018 |

**Best parameters**

**$\beta$ = 0.1**
**n_iter = 24**

**Patch 2**

Coordinates:
x1=53450
x2=54550
y1=8360
y2=9460

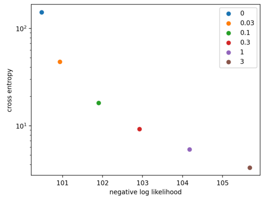


| beta | n_iter | LL | CE | ratio_CE_ lower_LL | CE_drop |
|---|---|---|---|---|---|
| 0 | 21 | -100.48 | 146.90 | 0.000 | 1.000 |
| 0.03 | 21 | -100.93 | 45.59 | 0.005 | 0.310 |
| **0.1** | **23** | **-101.90** | **17.21** | **0.014** | **0.117** |
| 0.3 | 26 | -102.92 | 9.25 | 0.024 | 0.063 |
| 1 | 32 | -104.18 | 5.73 | 0.037 | 0.039 |
| 3 | 50 | -105.68 | 3.72 | 0.052 | 0.025 |

**$\beta$ = 0.1**
**n_iter = 23**

**Patch 3**

Coordinates:
x1=55200
x2=56300
y1=7960
y2=9060

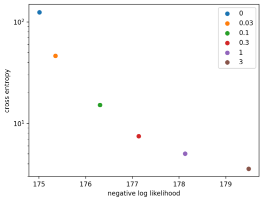


| beta | n_iter | LL | CE | ratio_CE_ lower_LL | CE_drop |
|---|---|---|---|---|---|
| 0 | 21 | -175.01 | 124.65 | 0.000 | 1.000 |
| 0.03 | 20 | -175.35 | 46.46 | 0.002 | 0.373 |
| **0.1** | **22** | **-176.31** | **15.16** | **0.007** | **0.122** |
| 0.3 | 25 | -177.14 | 7.49 | 0.012 | 0.060 |
| 1 | 28 | -178.13 | 5.03 | 0.018 | 0.040 |
| 3 | 39 | -179.49 | 3.57 | 0.026 | 0.029 |

**$\beta$ = 0.1**
**n_iter = 22**

**Patch 4**

Coordinates:
x1=56250
x2=57350
y1=8250
y2=9350

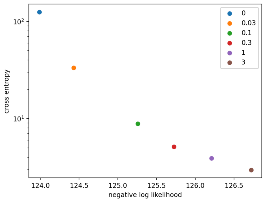


| beta | n_iter | LL | CE | ratio_CE_ lower_LL | CE_drop |
|---|---|---|---|---|---|
| 0 | 20 | -123.99 | 124.98 | 0.000 | 1.000 |
| 0.03 | 20 | -124.43 | 33.25 | 0.004 | 0.266 |
| **0.1** | **23** | **-125.26** | **8.86** | **0.010** | **0.071** |
| 0.3 | 26 | -125.72 | 5.11 | 0.014 | 0.041 |
| 1 | 30 | -126.21 | 3.89 | 0.018 | 0.031 |
| 3 | 40 | -126.72 | 2.96 | 0.022 | 0.024 |

**$\beta$ = 0.1**
**n_iter = 23**

**Patch 5**

Coordinates:
x1=45450
x2=46550
y1=4100
y2=5200

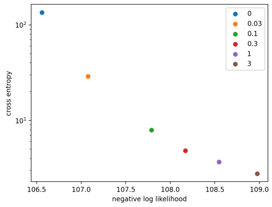


| beta | n_iter | LL | CE | ratio_CE_ lower_LL | CE_drop |
|---|---|---|---|---|---|
| 0 | 20 | -106.56 | 134.84 | 0.000 | 1.000 |
| 0.03 | 20 | -107.07 | 28.98 | 0.005 | 0.215 |
| **0.1** | **23** | **-107.79** | **7.93** | **0.012** | **0.059** |
| 0.3 | 27 | -108.17 | 4.84 | 0.015 | 0.036 |
| 1 | 31 | -108.55 | 3.68 | 0.019 | 0.027 |
| 3 | 42 | -108.97 | 2.78 | 0.023 | 0.021 |

**$\beta$ = 0.1,**
**n_iter = 23**

**Patch 6**

Coordinates:
x1=62800
x2=63900
y1=11000
y2=12100

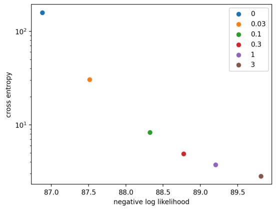


| beta | n_iter | LL | CE | ratio_CE_ lower_LL | CE_drop |
|---|---|---|---|---|---|
| 0 | 21 | -86.88 | 158.67 | 0.000 | 1.000 |
| 0.03 | 21 | -87.51 | 30.45 | 0.007 | 0.192 |
| **0.1** | **24** | **-88.32** | **8.31** | **0.017** | **0.052** |
| 0.3 | 27 | -88.77 | 4.92 | 0.022 | 0.031 |
| 1 | 31 | -89.21 | 3.75 | 0.027 | 0.024 |
| 3 | 47 | -89.81 | 2.84 | 0.034 | 0.018 |

**$\beta$ = 0.1**
**n_iter = 24**

**Supplementary Fig.15. Hyperparameter tuning**. Left: tradeoff curve between cross entropy and log likelihood under different neighborhood parameter beta. Higher beta puts more focus on optimizing cross entropy, leading to smoother but underfitted results with lower log likelihood. Right: table of training details for each beta value, where the iteration steps were selected from automatic stopping criterion under the current beta value. The n_iter is the number of iterations when training is converged; LL: log likelihood; CE: the cross entropy; ratio_LL_drop: relative drop of LL compared to no-neighbor model; CE_drop: the ratio of cross entropy between current beta and no-neighbor model. Top to bottom, hyper parameter tuning for 6 different patches cropped from the human colon cancer sample P2.

**Supplementary Fig.16.**

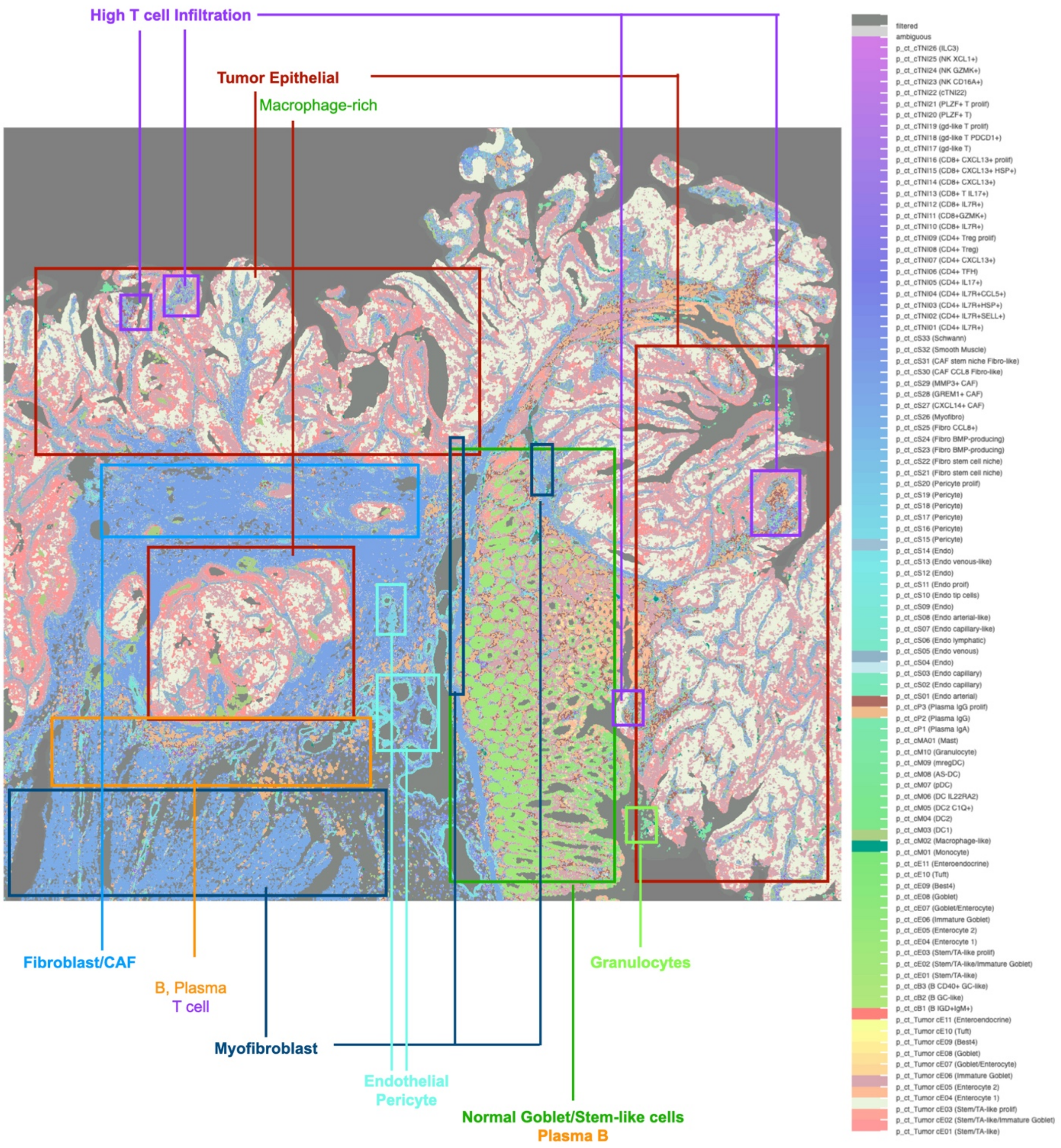


**Supplementary Fig.16. Overall tumor architecture based on STHD spot cell types.** Manual labeling of various global regions highlighted by STHD cell type labels.

**Supplementary Fig.17.**

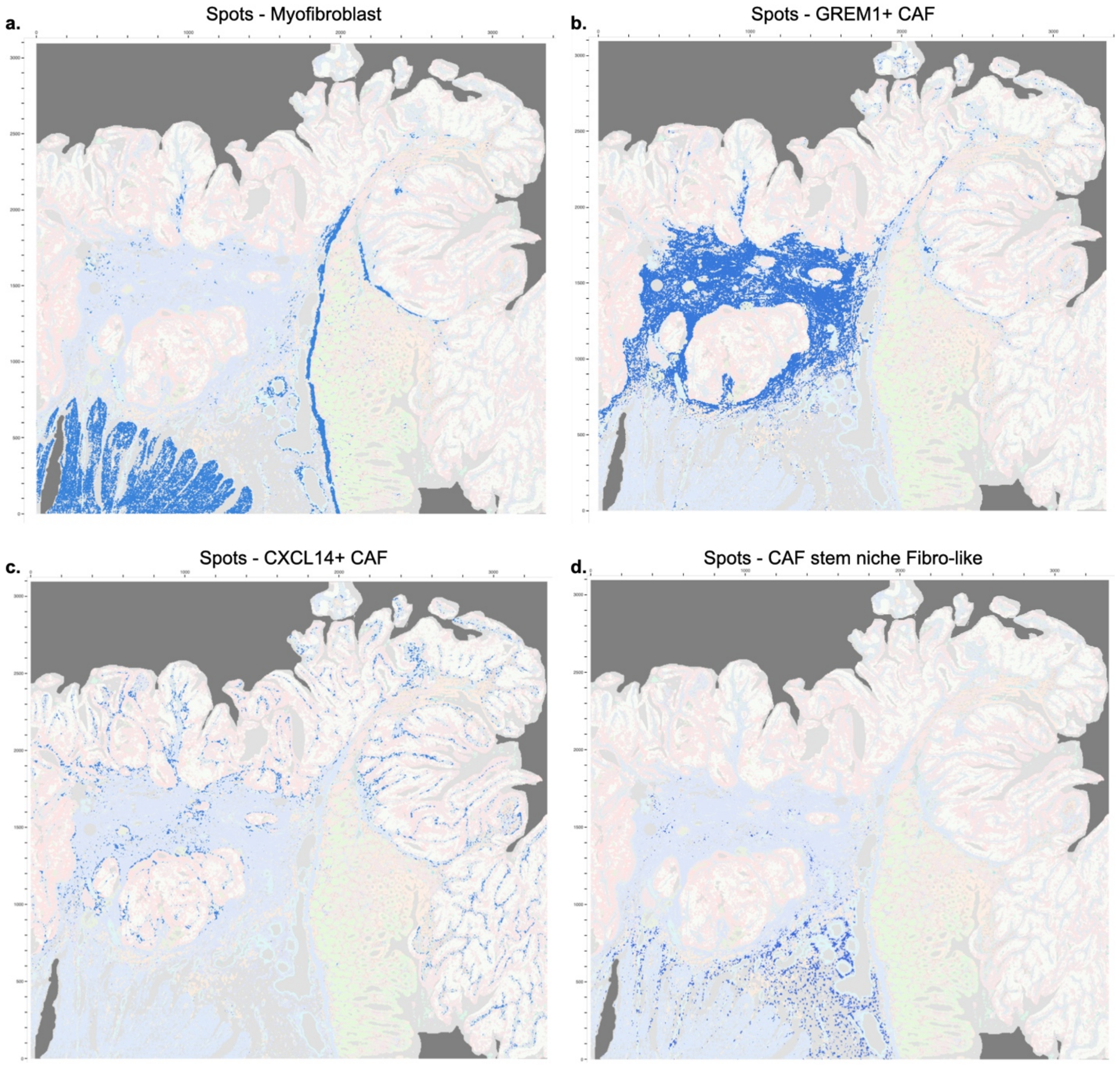


**Supplementary Fig.17. Example of spatially organized myofibroblast and fibroblast types.** **a.** Spots labeled as myofibroblasts. **b.** Spots labeled as GREM1+ CAF. **c.** Spots labeled as CXCL14+ CAF. **d.** Spots labeled as fibroblast-like stem niche CAFs. CAF, cancer associated fibroblasts.

**Supplementary Fig.18.**

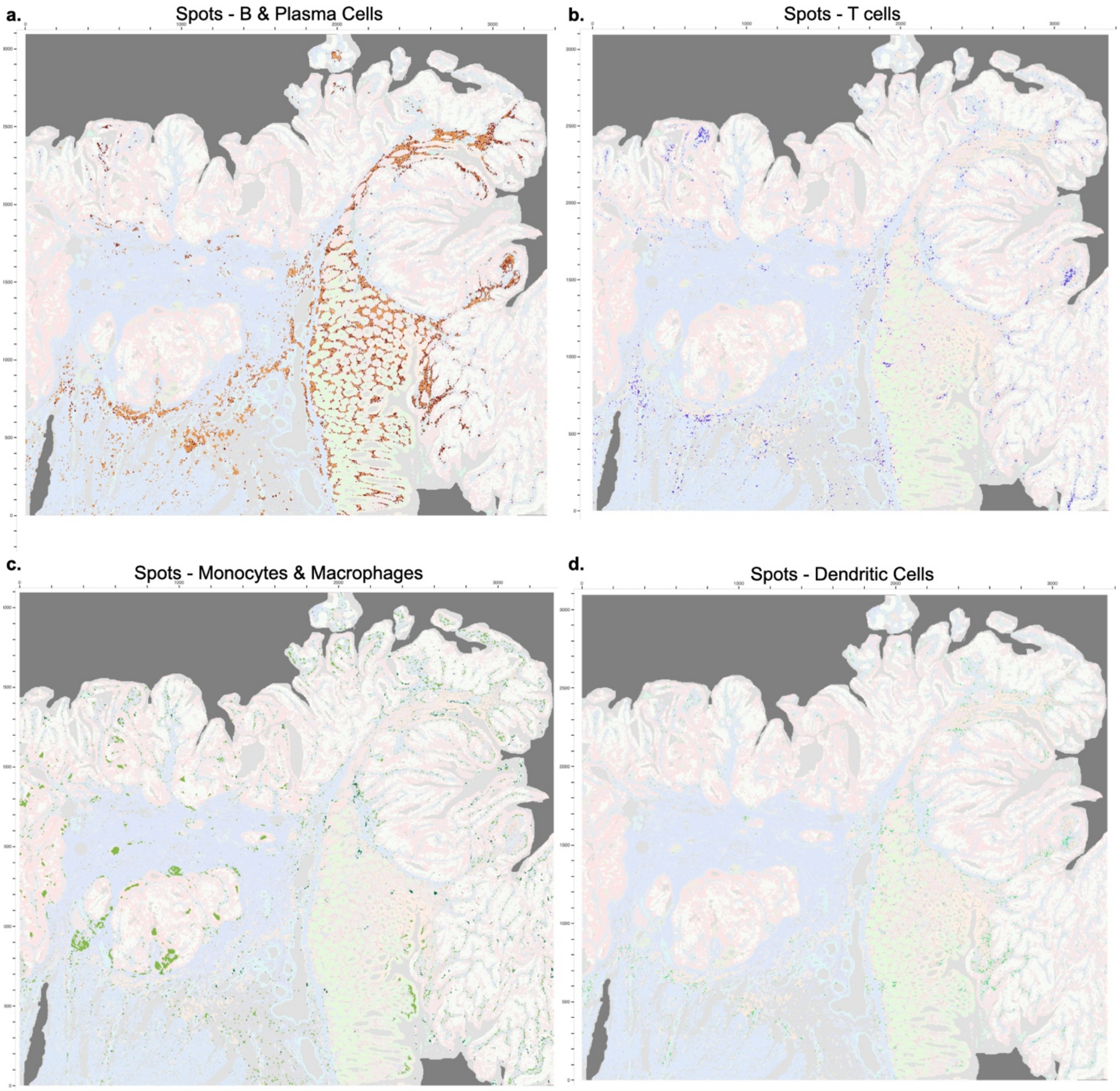


**Supplementary Fig.18. Spatially organized cell types of immune lineage. a.** Spots labeled as B cells or plasma cells, mostly interspaced in the gland region. **b.** Spots labeled as T cells including CD4T, CD8T, NK, and other T cells. **c.** Spots labeled as monocytes and macrophages. **d.** Spots labeled as dendritic cells.

**Supplementary Fig.19.**

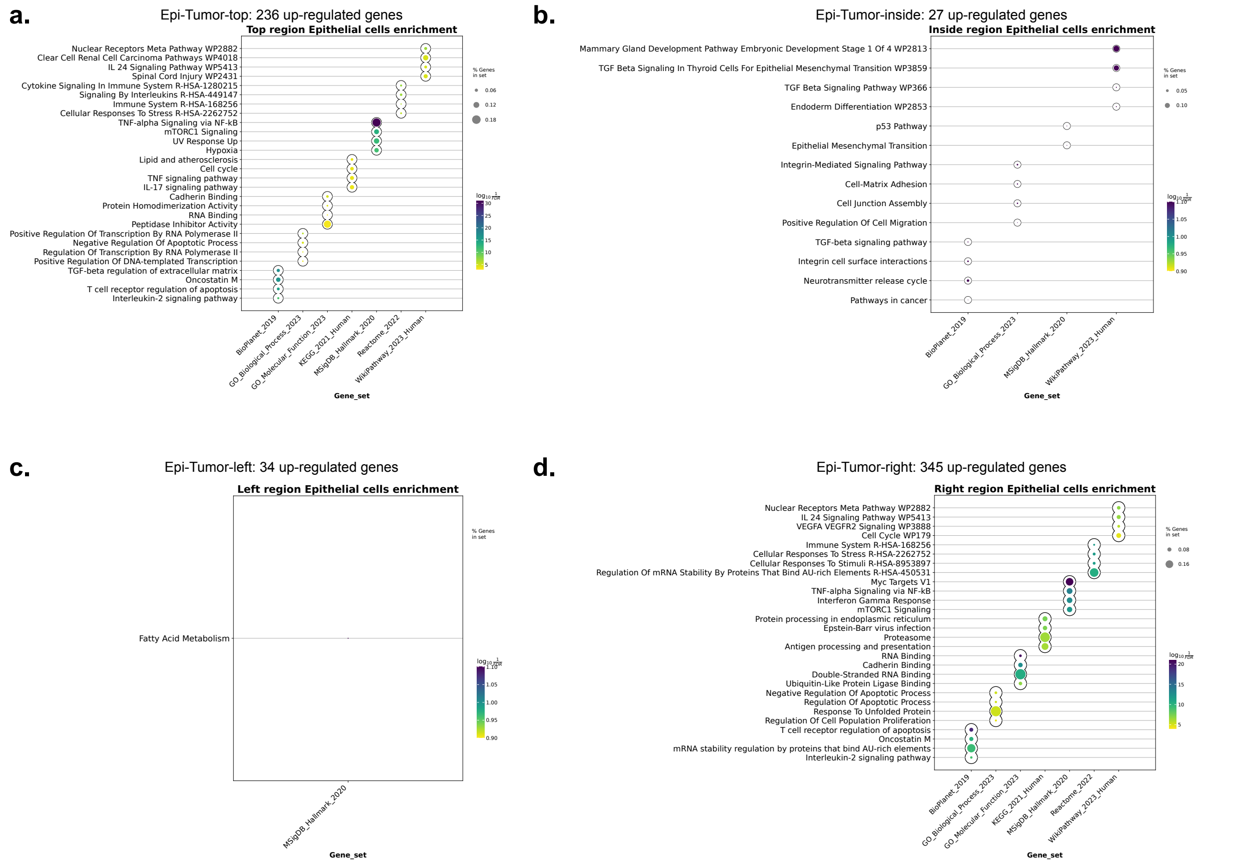


**Supplementary Fig.19. Pathway enrichment of epithelial-specific up-regulated genes in different spatial regions. a.** Epithelial-specific pathways in region Epi-Tumor-top. **b.** Epithelial-specific pathways in region Epi-Tumor-inside. **c.** Epithelial-specific pathways in region Epi-Tumor-left. **d.** Epithelial-specific pathways in region Epi-Tumor-right.

**Supplementary Fig.20.**

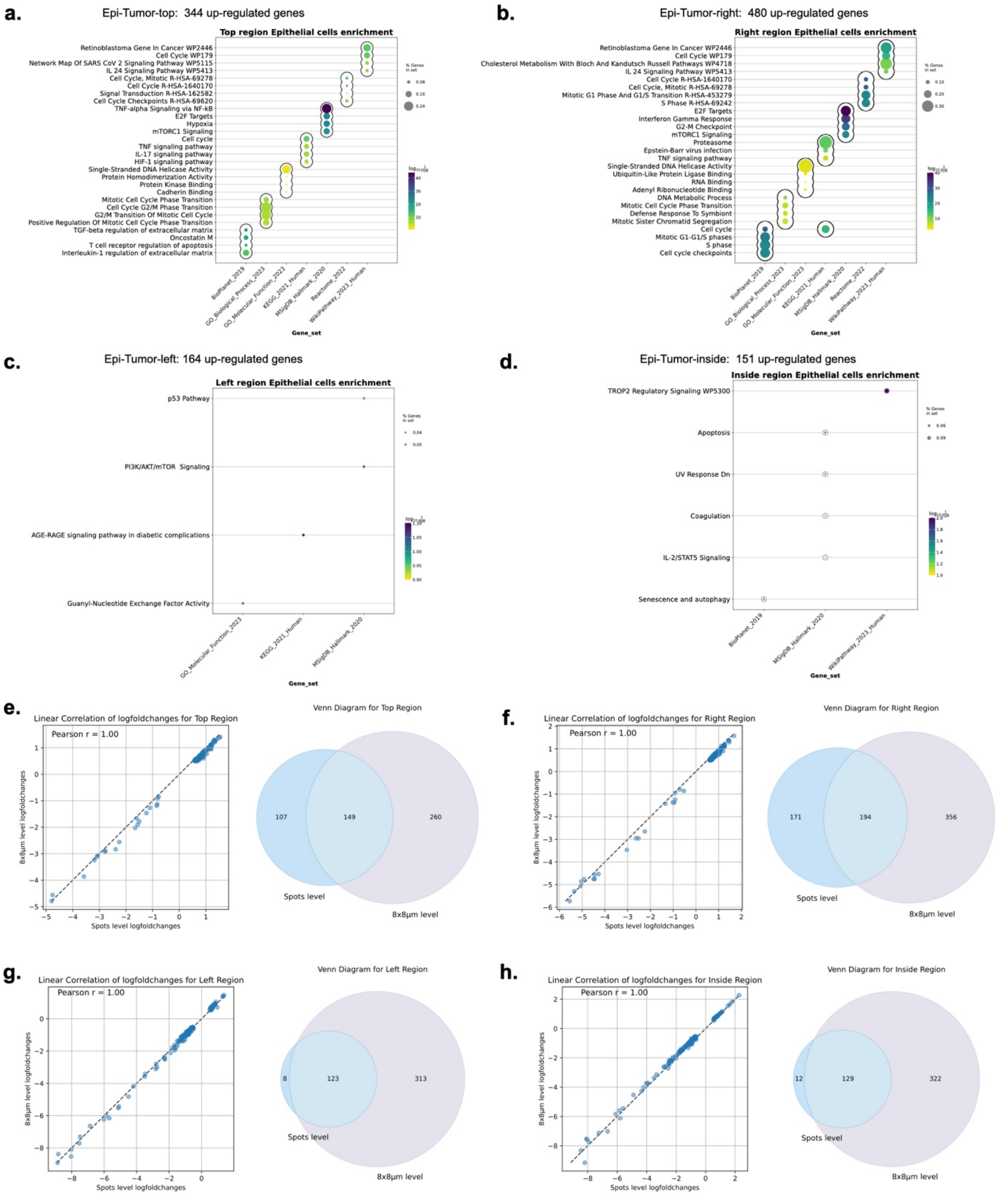


**Supplementary Fig.20. Enriched pathways specific to epithelial cell types across spatial regions**. Results of epithelial-specific and region-specific differentially expressed genes at STHD-guided bin level. **a-d.** Pathway enrichment performed on differentially upregulated genes specific to epithelial cell types, using STHD-guided bins of size 4x4 spots (8x8um). The regions are Epi-Tumor-top, Epi-Tumor-right, Epi-Tumor-left, Epi-Tumor-inside. **e-h.** Correlation between DE analyses between spot level and STHD-guided bin level. Left: Correlation of log fold changes between DE at spot-level or STHD-guided bin level, for epithelial-specific genes. Right: Venn diagram of DE genes obtained from spot-level versus STHD-guided bin level. DE, differential expression.

**Supplementary Fig.21.**

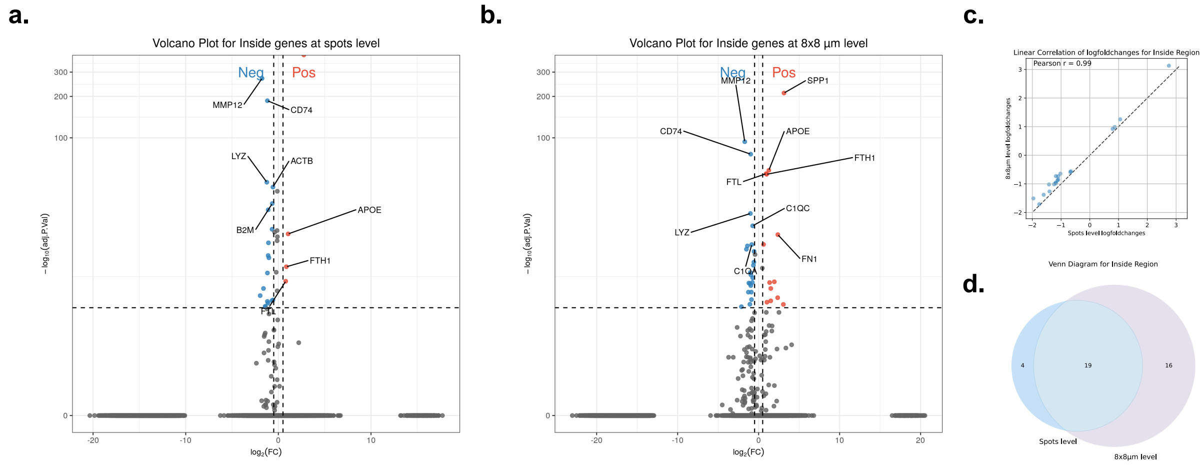


**Supplementary Fig.21. Macrophage-specific differential genes in tumor inside region. a.** macrophage-specific differentially expressed genes using STHD spot labels. Pos, differentially upregulated genes for macrophage spots in Tumor-Epi-inside region. Neg, depleted genes for macrophage spots in Tumor-Epi-inside region**. b.** macrophage-specific differentially expressed genes using STHD-guided bins in size of 4x4 spots. Pos, differentially upregulated genes for macrophage bins in Tumor-Epi-inside region. Neg, depleted genes for macrophage bins in Tumor-Epi-inside region**. c.** correlation of log fold change between spot-level differential analyses and bin-level differential analyses. **d.** Number of significantly differential genes overlapping spot-level and bin-level analyses.

**Supplementary Fig.22.**

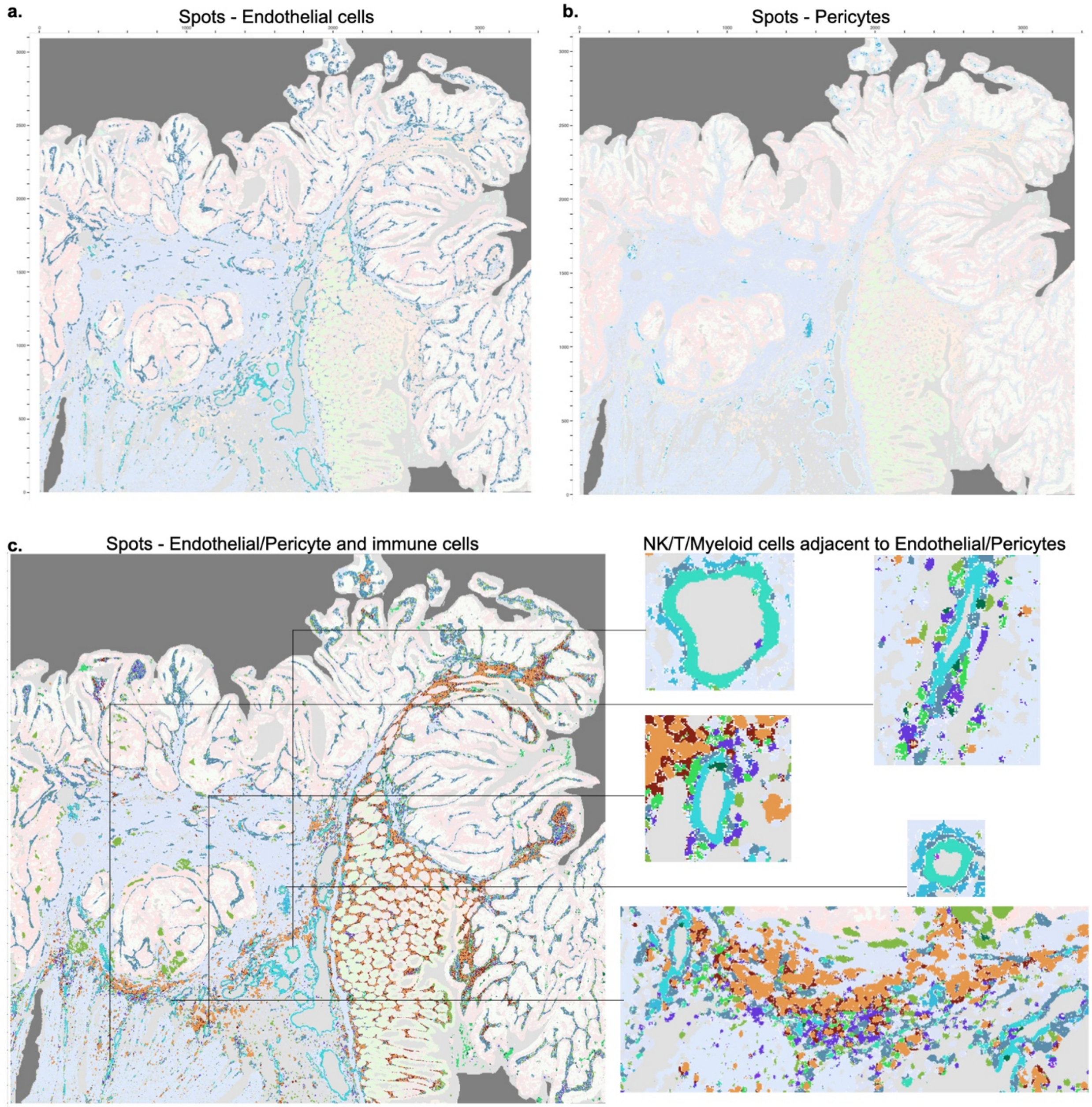


**Supplementary Fig.22. Endothelial cells and pericytes with vascular structures with adjacent infiltrating immune cells. a.** Spots labeled as endothelial cells. **b.** Spots labeled as pericytes usually surrounded around endothelial cells. **c.** Coloring immune cells with endothelial and pericytes. Immune cells include T cells in purple, monocyte/macrophages in green, and B/Plasma cells in orange and brown. Several locations are enlarged for immune cells infiltrating through endothelial vascular structures.

**Supplementary Fig.23.**

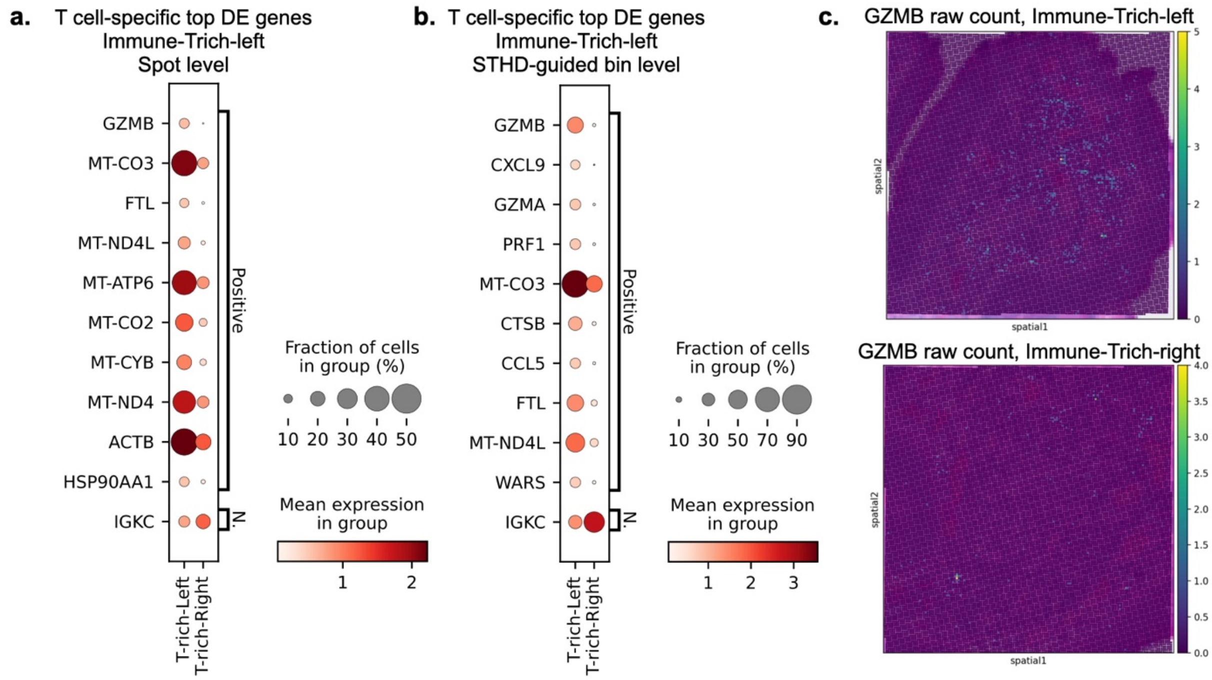


**Supplementary Fig.23. T cell-specific differentially expressed genes in two immune-rich regions. a.** At spot level, Genes significantly differentially expressed between Immune-Trich-left region and Immune-Trich-right region are shown. Top ten genes significantly higher in Immune-Trich-left is shown as Positive genes, where the only gene significant lower is IGKC. Genes are ranked by log fold change between the two immune-rich regions. **b.** Similar dot plot at STHD-guided bin level of size 4x4 spots. **c.** The raw expression counts of GZMB at spot level from the two regions from the colon cancer sample P2.

**Supplementary Fig.24.**

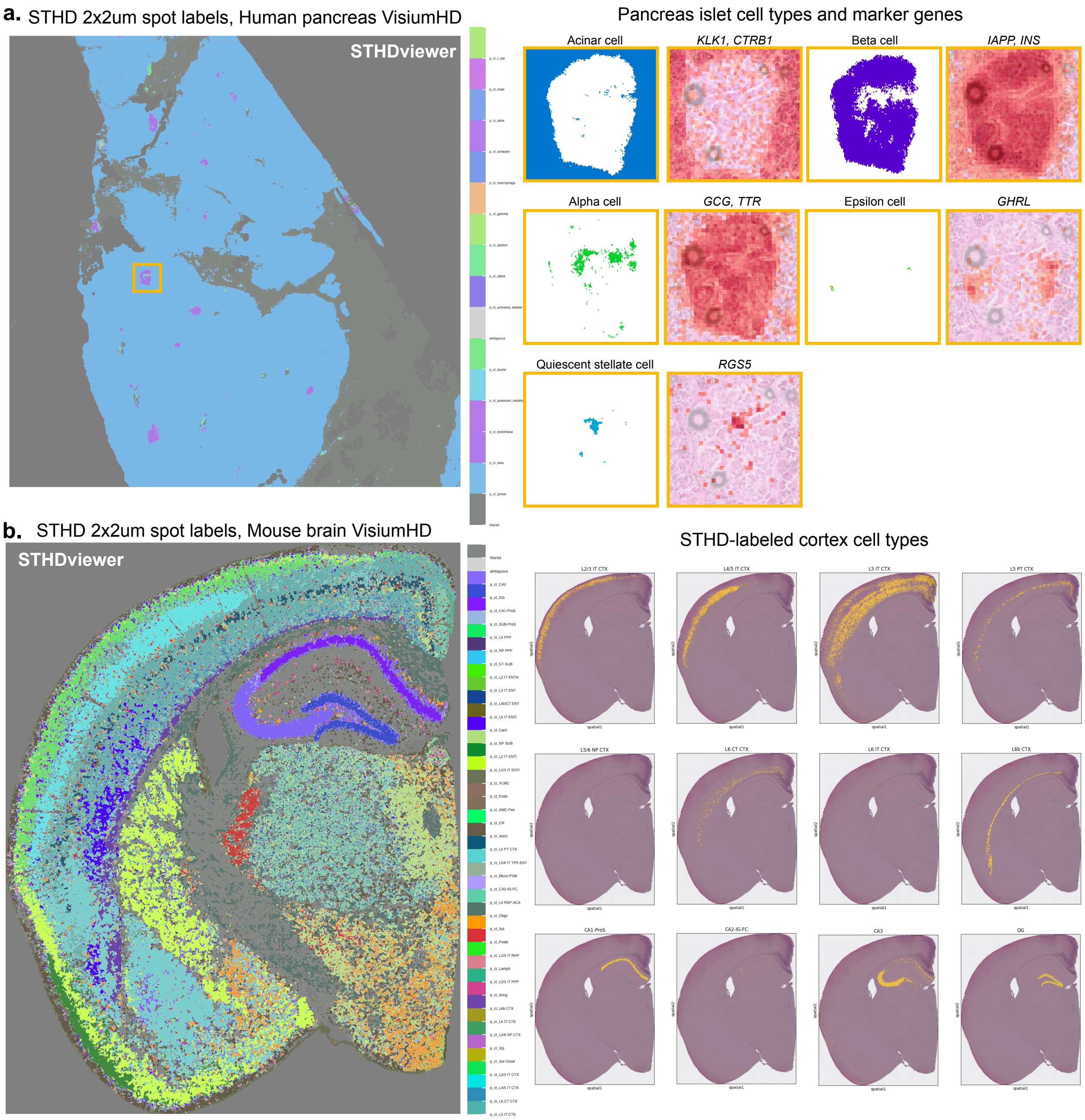


**Supplementary Fig.24. STHD spot labels for VisiumHD samples of human pancreas and mouse brain tissues. a.** Left: STHD labels of 2x2um spots in a human pancreas VisiumHD sample using an inDrop single-cell RNA reference. Right: a zoomed-in islet region, showing paired plots of various pancreatic cell types and marker gene expression visualized in Loupe browser with 8x8um bin resolution. **b.** Left: STHD labels of 2x2um spots in a mouse brain VisiumHD sample using Allen Brain Atlas reference data and annotation. Right: spots in cortex brain layers and hippocampal regions are highlighted in yellow.
