## Supplementary File 1 for "STHD: probabilistic cell typing of single Spots in whole Transcriptome spatial data with High Definition"

Yi Zhang, PhD[1]

[1]Department of Neurosurgery, Department of Biostatistics and Bioinformatics, Duke Cancer Institute, Duke University

### 1 Abstract

Recent spatial transcriptomics (ST) technologies have enabled single- and sub-cellular resolution profiling of gene expression across the whole transcriptome. However, the transition to high-definition ST significantly increased data sparsity and dimensionality, posing computational challenges in identifying cell types, deciphering neighborhood structure, and detecting differential expression - all are crucial steps to study normal and disease ST samples. Here we present STHD, a novel machine learning method for probabilistic cell typing of single spots in whole-transcriptome, high-resolution ST data. Unlike the current binning-aggregation-deconvolution strategy, STHD directly models gene expression at single-spot level to infer cell type identities without cell segmentation or spot aggregation. STHD addresses sparsity by modeling count statistics, incorporating neighbor similarities, and leveraging reference single-cell RNA-seq data. We show in VisiumHD data that STHD accurately predicts cell type identities at single-spot level, which achieves precise segmentation of both global tissue architecture and local multicellular neighborhoods. The high-resolution labels facilitate various downstream analyses, including cell type-stratified bin aggregation, spatial compositional comparisons, and cell type-specific differential expression analyses. Moreover, STHD labels further reveal frontlines of inter-cell type interactions at immune hubs in cancer samples. STHD is scalable and generalizable across diverse samples, tissues, and diseases, facilitating genome-wide analyses in various spatial organization contexts. Overall, computational modeling of individual spots with STHD facilitates discoveries in cellular interactions and molecular mechanisms in whole-genome spatial technologies with high resolution.

### 2 STHD model

STHD is a machine learning method that models raw counts of each gene at each spatial spot while incorporating neighbor spot information. The STHD loss function contains two parts, negative log likelihood of observed counts $-LL$, and cross entropy between neighboring spots $CE$.

$$\mathcal{L} = -LL + \beta CE$$

Let the normalized average gene expression level for gene $g$ within the latent cell type $t$ is $\lambda_g^t$. The average normalized gene expression $\lambda_g^t$ can be estimated from reference single-cell RNA-seq dataset where annotation of cell type $t$ is available. Before normalizing genes, selection of a set of cell type-informative genes is recommended. For each cell type, average normalized gene expression profiles are estimated with $\lambda_g^t = \frac{n_g^t}{\sum_g n_g^t}$ so that $\sum_g \lambda_g^t = 1$.

In the spatial data, suppose the total count depth in spot $a$ is $d_a$, then the gene count $n_g^a$ at spot $a$ given cell type $t$ follows:

$$n_g^a|t \sim Poisson(d_a \lambda_g^t)$$

The probability to obtain measurement $n_g^a$ with latent cell type $t \in T$ is:

$$P(n_g^a) = \sum_{t \in T} P(n_g^a|t) P_a(t)$$

Thus,

$$P(n_g^a|t) = \frac{(d_a \lambda_g^t)^{n_g^a} e^{-d_a \lambda_g^t}}{n_g^a!}$$

Log likelihood:

$$\begin{aligned}
LL &= log \prod_a \prod_g P(n_g^a) \\
&= \sum_a \sum_g log P(n_g^a) \\
&= \sum_a \sum_g log E_{P_a(t)}[P(n_g^a|t)] \\
&\geq \sum_a \sum_g E_{P_a(t)}[log P(n_g^a|t)] \\
&= \sum_a \sum_t P_a(t) \sum_g \left( [n_g^a log(d_a \lambda_g^t) - d_a \lambda_g^t] - log(n_g^a!) \right) \\
&= \sum_a \sum_t P_a(t) \sum_g \left( n_g^a log(d_a \lambda_g^t) - d_a \lambda_g^t \right) + const
\end{aligned} \tag{1}$$

We wish to estimate $P_a(t)$. Since spots are mostly subcellular and nearby spots tend to have similar cell types, and also because measurements at spots are sparse, we add regularization to the STHD loss $\mathcal{L}$ considering similarity between $P_a(t)$ and $P_{a'}(t)$ for spot $a$ and a neighboring spot $a'$ in the set of defined nearest neighbors $NN(a)$. Cross Entropy loss is thus:

$$CE = -\sum_a \sum_{a' \in NN(a)} \sum_t \left( P_a(t) log P_{a'}(t) \right) \tag{2}$$

For the full cell type set $T$, one constraint is $\sum_{t \in T} P_a(t) = 1$. Let

$$P_a(t) \equiv \frac{e^{w_a^t}}{\sum_{s \in T} e^{w_a^s}} \tag{3}$$

Thus, we optimize $P_a(t)$ or $w_a^t$ to minimize total loss $\mathcal{L}$:

$$\begin{aligned}
\mathcal{L} &= -LL + \beta CE \\
&= -\sum_a \sum_t P_a(t) \sum_g \left( n_g^a log(d_a \lambda_g^t) - d_a \lambda_g^t \right) - \beta \sum_a \sum_{a' \in NN(a)} \sum_t \left( P_a(t) log P_{a'}(t) \right)
\end{aligned} \tag{4}$$

Annotation:
$a \in A$: A set of HD spots in the region of interest
$g \in G$: Genes used for cell typing
$t \in T$: Set of cell types
$a' \in NN(a)$: Neighboring spots of $a$

### 2.1 Derivative of log likelihood

For fast STHD implementation, we here compute explicit form of derivative of loss function $\mathcal{L}$ for efficient optimization. For a particular pair of spot $\tilde{a}$ and cell type identity $\tilde{t}$, we first derive:

$$\frac{\partial P_a(t)}{\partial w_{\tilde{a}}^{\tilde{t}}} = \frac{\partial}{\partial w_{\tilde{a}}^{\tilde{t}}}(\frac{e^{w_a^t}}{\sum_{s\in T} e^{w_a^s}})$$

Let

$$\pi_a \equiv \sum_{s\in T} e^{w_a^s}$$

If $a \neq \tilde{a}$:

$$\frac{\partial P_a(t)}{\partial w_{\tilde{a}}^{\tilde{t}}} = 0$$

If $a = \tilde{a}$ and $t \neq \tilde{t}$:

$$\begin{aligned}
\frac{\partial P_a(t)}{\partial w_{\tilde{a}}^{\tilde{t}}} &= \frac{\partial}{\partial w_{\tilde{a}}^{\tilde{t}}}(\frac{e^{w_a^t}}{\sum_{s\in T} e^{w_a^s}}) \\
&= \frac{\partial}{\partial w_{a}^{\tilde{t}}}(\frac{e^{w_a^t}}{\sum_{s\in T} e^{w_a^s}}) \\
&= e^{w_a^t}\frac{\partial}{\partial w_{a}^{\tilde{t}}}(\frac{1}{\sum_{s} e^{w_a^s}}) \\
&= -e^{w_a^t}(\frac{1}{\sum_{s} e^{w_a^s}})^2 * e^{w_a^{\tilde{t}}} \\
&= -\frac{e^{w_a^t} e^{w_a^{\tilde{t}}}}{\pi_a^2} \\
&= -\frac{e^{w_{\tilde{a}}^t} e^{w_{\tilde{a}}^{\tilde{t}}}}{\pi_{\tilde{a}}^2} \\
&= -P_{\tilde{a}}(t)P_{\tilde{a}}(\tilde{t})
\end{aligned}$$

If $a = \tilde{a}$ and $t = \tilde{t}$:

$$
\begin{aligned}
\frac{\partial P_a(t)}{\partial w_{\tilde{a}}^{\tilde{t}}} &= \frac{\partial}{\partial w_{\tilde{a}}^{\tilde{t}}}(\frac{e^{w_a^t}}{\sum_{s\in T} e^{w_a^s}}) \\
&= \frac{\partial}{\partial w_a^t}(\frac{e^{w_a^t}}{\sum_{s\in T} e^{w_a^s}}) \\
&= e^{w_a^t}\frac{1}{\sum_s e^{w_a^s}} + e^{w_a^t}\frac{\partial}{\partial e^{w_a^t}}(\frac{1}{\sum_s e^{w_a^s}}) \\
&= \frac{e^{w_a^t}}{\pi_a} - (\frac{e^{w_a^t}}{\pi_a})^2 \\
&= \frac{e^{w_{\tilde{a}}^{\tilde{t}}}}{\pi_{\tilde{a}}} - (\frac{e^{w_{\tilde{a}}^{\tilde{t}}}}{\pi_{\tilde{a}}})^2 \\
&= P_{\tilde{a}}(\tilde{t}) - P_{\tilde{a}}(\tilde{t})^2
\end{aligned}
$$

Let

$$
F(a,t) = \sum_g \Big(n_g^a log(d_a\lambda_g^t) - d_a\lambda_g^t\Big)
$$

Thus, derivative of $LL$ :

$$
\begin{aligned}
\frac{\partial LL}{\partial w_{\tilde{a}}^{\tilde{t}}} &= \sum_a\sum_t \frac{\partial P_a(t)}{\partial w_{\tilde{a}}^{\tilde{t}}}F(a,t) \\
&= \sum_{a\neq\tilde{a}}\sum_t \frac{\partial P_a(t)}{\partial w_{\tilde{a}}^{\tilde{t}}}F(a,t) + \sum_{a=\tilde{a}}\sum_{t\neq\tilde{t}} \frac{\partial P_{\tilde{a}}(t)}{\partial w_{\tilde{a}}^{\tilde{t}}}F(\tilde{a},t) + \sum_{a=\tilde{a}}\sum_{t=\tilde{t}} \frac{\partial P_{\tilde{a}}(\tilde{t})}{\partial w_{\tilde{a}}^{\tilde{t}}}F(\tilde{a},\tilde{t}) \\
&= 0 - \sum_{t\neq\tilde{t}} P_{\tilde{a}}(t)P_{\tilde{a}}(\tilde{t})F(\tilde{a},t) + \Big(P_{\tilde{a}}(\tilde{t}) - P_{\tilde{a}}(\tilde{t})^2\Big)F(\tilde{a},\tilde{t}) \\
&= P_{\tilde{a}}(\tilde{t})F(\tilde{a},\tilde{t}) - \sum_t P_{\tilde{a}}(t)P_{\tilde{a}}(\tilde{t})F(\tilde{a},t)
\end{aligned}
$$

### 2.2 Derivative of Cross Entropy Loss

For spot $a$, we pre-define its nearest neighbor set $NN(a)$, marking one of them $a'$. Cross Entropy loss:

$$
CE = -\sum_a \sum_{a'\in NN(a)} \sum_t \Big(P_a(t) log P_{a'}(t)\Big)
$$

For a particular pair of spot $\tilde{a}$ and cell type identity $\tilde{t}$, the derivative of cross entropy loss:

$$
\begin{aligned}
\frac{\partial CE}{\partial w_{\tilde{a}}^{\tilde{t}}} &= -\sum_{a}\sum_{a'\in NN(a)}\sum_{t}\Big(\frac{\partial P_a(t)}{\partial w_{\tilde{a}}^{\tilde{t}}} logP_{a'}(t) + \frac{P_a(t)}{P_{a'}(t)}\frac{\partial P_{a'}(t)}{\partial w_{\tilde{a}}^{\tilde{t}}}\Big) \\
&= -\sum_{a=\tilde{a}}\sum_{a'\in NN(a)}\sum_{t}\Big(\frac{\partial P_{\tilde{a}}(t)}{\partial w_{\tilde{a}}^{\tilde{t}}} logP_{a'}(t) + \frac{P_{\tilde{a}}(t)}{P_{a'}(t)}\frac{\partial P_{a'}(t)}{\partial w_{\tilde{a}}^{\tilde{t}}}\Big) \quad (5)\\
&\quad -\sum_{a\neq\tilde{a}}\sum_{a'\in NN(a),a'=\tilde{a}}\sum_{t}\Big(\frac{\partial P_a(t)}{\partial w_{\tilde{a}}^{\tilde{t}}} logP_{\tilde{a}}(t) + \frac{P_a(t)}{P_{a'}(t)}\frac{\partial P_{\tilde{a}}(t)}{\partial w_{\tilde{a}}^{\tilde{t}}}\Big) \quad (6)\\
&\quad -\sum_{a\neq\tilde{a}}\sum_{a'\in NN(a),a'\neq\tilde{a}}\sum_{t}\Big(\frac{\partial P_a(t)}{\partial w_{\tilde{a}}^{\tilde{t}}} logP_{a'}(t) + \frac{P_a(t)}{P_{a'}(t)}\frac{\partial P_{a'}(t)}{\partial w_{\tilde{a}}^{\tilde{t}}}\Big) \quad (7)\\
&= (5) + (6) + (7)
\end{aligned}
$$

From the previous section, we know that in scenarios (5) and (7), $a' \neq \tilde{a}$, so $\frac{\partial P_{a'}(t)}{\partial w_{\tilde{a}}^{\tilde{t}}} = 0$ . In scenario (7), $a \neq \tilde{a}$, thus $\frac{\partial P_a(t)}{\partial w_{\tilde{a}}^{\tilde{t}}} = 0$ , and (7) $= 0$. Also, $a' \neq a$ because they are neighbors.

$$
\begin{aligned}
(5) &= -\sum_{a=\tilde{a}}\sum_{a'\in NN(a)}\sum_{t}\Big(\frac{\partial P_{\tilde{a}}(t)}{\partial w_{\tilde{a}}^{\tilde{t}}} logP_{a'}(t)\Big) \\
&= -\sum_{a'\in NN(a)}\Big[\Big(P_{\tilde{a}}(\tilde{t}) - P_{\tilde{a}}(\tilde{t})^2\Big) logP_{a'}(\tilde{t}) + \sum_{t\neq\tilde{t}}\Big(-P_{\tilde{a}}(t)P_{\tilde{a}(\tilde{t})}\Big) logP_{a'}(t)\Big] \\
&= -P_{\tilde{a}}(\tilde{t})\sum_{a'\in NN(\tilde{a})}\Big(logP_{a'}(\tilde{t}) - \sum_{t} P_{\tilde{a}}(t) logP_{a'}(t)\Big)
\end{aligned}
$$

For scenario (6) , $a \neq \tilde{a}$, thus $\frac{\partial P_a(t)}{\partial w_{\tilde{a}}^{\tilde{t}}} = 0$.

$$
\begin{aligned}
(6) &= -\sum_{a\neq\tilde{a}}\mathbb{I}_{\tilde{a}\in NN(a)}\Big(\sum_{t=\tilde{t}}\frac{P_a(t)}{P_{\tilde{a}}(t)}\frac{\partial P_{\tilde{a}}(\tilde{t})}{\partial w_{\tilde{a}}^{\tilde{t}}} + \sum_{t\neq t'}\frac{P_a(t)}{P_{\tilde{a}(t)}}\frac{\partial P_{\tilde{a}}(t)}{\partial w_{\tilde{a}}^{\tilde{t}}}\Big) \\
&= -\sum_{a\neq\tilde{a}}\mathbb{I}_{\tilde{a}\in NN(a)}\frac{P_a(t)}{P_{\tilde{a}}(t)}\Big(P_{\tilde{a}}(\tilde{t}) - P_{\tilde{a}}(\tilde{t})^2\Big) + \sum_{t\neq t'}\frac{P_a(t)}{P_{\tilde{a}}(t)}\Big(-P_{\tilde{a}}(t)P_{\tilde{a}}(\tilde{t})\Big) \\
&= -\sum_{a\neq\tilde{a}}\mathbb{I}_{\tilde{a}\in NN(a)}\Big(P_a(\tilde{t}) - \sum_{t} P_a(t)P_{\tilde{a}}(\tilde{t})\Big) \\
&= -\sum_{a\neq\tilde{a}}\mathbb{I}_{\tilde{a}\in NN(a)}\Big(P_a(\tilde{t}) - P_{\tilde{a}}(\tilde{t})\Big)
\end{aligned}
$$

$$
\frac{\partial CE}{\partial w_{\tilde{a}}^{\tilde{t}}} = -P_{\tilde{a}}(\tilde{t})\sum_{a'\in NN(\tilde{a})}\Big(logP_{a'}(\tilde{t}) - \sum_{t} P_{\tilde{a}}(t) logP_{a'}(t)\Big) - \sum_{a\neq\tilde{a}}\mathbb{I}_{\tilde{a}\in NN(a)}\Big(P_a(\tilde{t}) - P_{\tilde{a}}(\tilde{t})\Big) \quad (8)
$$

#### 2.3 Derivative of STHD loss function

Thus, derivative of loss function is

$$
\begin{aligned}
\frac{\partial \mathcal{L}}{\partial w_{\tilde{a}}^{\tilde{t}}} &= -\frac{\partial LL}{\partial w_{\tilde{a}}^{\tilde{t}}} + \beta \frac{\partial CE}{\partial w_{\tilde{a}}^{\tilde{t}}} \\
&= -P_{\tilde{a}}(\tilde{t}) F(\tilde{a}, \tilde{t}) + \sum_t P_{\tilde{a}}(t) P_{\tilde{a}}(\tilde{t}) F(\tilde{a}, t) \\
&- \beta \Big[ P_{\tilde{a}}(\tilde{t}) \sum_{a' \in NN(\tilde{a})} \Big( log P_{a'}(\tilde{t}) - \sum_t P_{\tilde{a}}(t) log P_{a'}(t) \Big) - \sum_{a \neq \tilde{a}} \mathbb{I}_{\tilde{a} \in NN(a)} \Big( P_a(\tilde{t}) - P_{\tilde{a}}(\tilde{t}) \Big) \Big]
\end{aligned}
$$

Where

$$F(a, t) = \sum_g \Big( n_g^a log(d_a \lambda_g^t) - d_a \lambda_g^t \Big)$$

$$P_a(t) = \frac{e^{w_a^t}}{\sum_{s \in T} e^{w_a^s}}$$

STHD thus optimize $w_{\tilde{a}}^{\tilde{t}}$ and $P_{\tilde{a}}(\tilde{t})$ using gradient descent with Adam[1].
